# Stage-resolved dynamics of antibiotic resistance genes and mobile genetic elements in full-scale anaerobic digestion systems

**DOI:** 10.64898/2026.03.02.709106

**Authors:** He Sun, Sara Bergström Nilsson, Anna Schnürer

## Abstract

Anaerobic digestion (AD) has the potential to reduce antibiotic resistance genes (ARGs) and mobile genetic elements (MGEs) during waste treatment, yet evidence from full-scale plants remains limited and the relative importance of operational and physicochemical drivers is poorly resolved. Here, we profiled stage-resolved resistome and mobilome dynamics across substrates, digestates, and post-digestion storages from ten Swedish farm-scale AD plants (42 samples). High-throughput qPCR (ResistoMap) was combined with absolute 16S rRNA gene quantification and 16S rRNA gene amplicon sequencing to quantify ARG/MGE abundances and characterise bacterial community structure. ARGs and MGEs showed heterogeneous changes during digestion and subsequent storage, with both increases and decreases across plants. Non-metric multidimensional scaling (NMDS) suggested that higher pH and free ammonia were associated with decreasing trends in ARG and MGE relative abundance. Stage-resolved correlation networks revealed a dominant ARG-MGE association backbone that remained largely conserved from substrates to digestates and storage, whereas host-ARG and host-MGE associations were extensively restructured across processing stages. Collectively, these results show that stage-resolved, network-informed analyses offer a more informative framework than pooled, static approaches for evaluating resistance persistence and potential environmental risks associated with digestate application.

## 1. Introduction

Anaerobic digestion (AD) is a widely recognised cornerstone of circular agricultural systems, as it enables the conversion of organic manure and crop residues into renewable energy (biogas) and nutrient-rich digestates.^1^ Digestates derived from AD systems are widely applied as organic fertilisers due to their high availability of nitrogen, phosphorus, and potassium, as well as fertilisation efficiencies comparable to mineral fertilisers.^2^ In addition, biogas production contributes to climate mitigation by reducing greenhouse gas emissions and improving on-farm energy balance.^3^ As a result, the large-scale reuse of digestates on agricultural land has increased substantially across Europe.^4^ Given the central role of digestates in nutrient recycling within circular agriculture, ensuring their safe application at scale is essential to prevent the release of chemical or biological contaminants into the environment.

Among the potential risks associated with digestate application, growing attention has been directed toward the unintended dissemination of antibiotic resistance. Antibiotic resistance is widely recognized as a major global health challenge spanning human, animal, and environmental domains.^5,6^ Digestates originating from livestock manure, food waste, and other organic substrates have been shown to contain antibiotic resistance genes (ARGs) and mobile genetic elements (MGEs), which together determine both the persistence and mobility of resistance in environmental systems.^7–9^ While ARGs are largely introduced via incoming substrates rather than generated during digestion,^7,10^ their fate during AD remains a central concern for assessing downstream environmental risks.

Previous studies have reported both reductions and enrichments of ARGs during AD, with outcomes varying widely across systems.^10–14^ Such variability partly reflects gene-specific responses, whereby certain ARGs may be enriched even when the majority of resistance genes are reduced under the same digestion conditions. For instance, *tet*W, *erm*B and *erm*F were reported to increase in abundance during mesophilic AD of wastewater sludge, while six other ARGs were simultaneously reduced.^15^ Beyond gene-specific behaviour, the extent of ARG reduction during AD is strongly influenced by experimental design and biogas process parameters, including substrate composition, pre-treatment methods, operational temperature, retention time, organic loading rate, and downstream handling practices.^7,10^ In this context, several studies have demonstrated that thermophilic AD generally achieves greater ARG reduction than mesophilic AD,^11,16,17^ and two-stage (acidogenic-methanogenic) systems have been reported to outperform single-stage AD in ARG removal in some cases.^18,19^ In addition, high-solids AD was found to be more effective in ARG reduction than wet-type AD.^20,21^ Despite recognition of these influencing factors, systematic and process-oriented investigations remain scarce, particularly studies that identify which operational parameters most strongly affect resistance reduction across the biogas production chain. Consequently, although AD is often regarded as a promising strategy for mitigating antibiotic resistance, current knowledge is insufficient to define operational conditions that consistently minimise resistance risks under full-scale, real-world settings, as most existing studies are laboratory-based and focus on a limited number of ARGs.^14–16,21,22^

Beyond changes in ARG abundance during AD, increasing attention has been directed toward host–ARG and ARG–MGE associations, as ARGs can persist within microbial hosts via vertical gene transfer (VGT) and disseminate among bacteria through horizontal gene transfer (HGT) mediated by MGEs.^11,23^ Correlation-based network analyses have therefore been widely applied to infer host–ARG and ARG–MGE relationships, providing insights into potential resistance carriers and mobility pathways during AD.^21,24^ However, most existing studies have constructed such networks based on aggregated samples, by pooling substrates and digestates, likely due to limitations in sample size.^11,19,21,24^ As a result, such analyses predominantly provide static snapshots of ARG-MGE-host associations. Consequently, the redistribution of ARGs among microbial hosts across successive AD stages, as well as the persistence or disruption of underlying resistance–mobility structures, remains poorly understood. Moreover, associations inferred from pooled samples may not be consistently supported at individual AD stages, potentially constraining interpretations of ARG fate based on changes in the abundance of putative host taxa.

The aim of this study was therefore to investigate the dynamics of ARGs and MGEs across different stages of full-scale AD systems. Samples were collected from ten farm-scale biogas plants in Sweden, representing organic, conventional, and mixed organic-conventional farming practices. An integrated approach combining high-throughput quantitative PCR (qPCR) targeting ARGs and MGEs with 16S rRNA gene PCR and qPCR was applied to characterise the resistome, mobilome, and bacterial community structure and abundance. This framework was used to assess how key operational parameters and process configurations influence the reduction of antibiotic resistance, as well as the redistribution of ARGs among microbial hosts and MGEs along the AD process. Overall, the results provide a process-oriented basis for improving digestate management and mitigating the potential environmental dissemination of antibiotic resistance.

## 2. Materials and Methods

### 2.1 Samples and biogas plant operations

A total of 42 samples were collected from ten full-scale biogas plants distributed across Sweden. The samples encompassed a wide range of substrate types, including cow, pig, and chicken manure, slaughterhouse waste, beddings, and plant-derived materials, as well as the corresponding digestates and post-digestion storage samples (Table S1). The biogas plants were classified according to substrate origin into organic (n = 3), conventional (n = 3), and mixed conventional–organic systems (n = 4). Operating conditions varied substantially among the plants, with digester temperatures at the time of sampling ranging from 35 to 51 °C and hydraulic retention times (HRT) from 17 and 75 days. Additional operational parameters and sample characteristics, including pH, organic loading rate (OLR), ammonium nitrogen (NH_4_^+^– N), ammonia nitrogen (NH_3_–N), total solids (TS), volatile solids (VS), volatile fatty acids (VFAs) and digestate handling practices, are summarised in Table S2. All samples were stored at −20 °C prior to DNA extraction and physicochemical analyses.

### 2.2 Physicochemical analyses

TS and VS were determined in triplicate using standard methods.^25^ Concentrations of NH_4_^+^–N and VFAs, as well as pH, were measured following procedures described previously.^26^ NH_3_–N concentrations were calculated from temperature, pH, and NH_4_^+^–N according to a published approach.^27^

### 2.3 DNA extraction and molecular quantification

DNA was extracted in triplicate from all samples using the FastDNA Spin Kit for Soil (MP Biomedicals Europe) following the manufacturer’s instructions. Extracted DNA was purified using AMPure magnetic beads (Beckman Coulter) following the manufacturer’s protocol, and DNA concentrations were quantified using a Qubit fluorometer. Relative abundances of ARGs and MGEs, normalised to 16S rRNA gene copy numbers, were determined using a high-throughput qPCR platform (ResistoMap, https://www.resistomap.com). Absolute abundances of total bacteria were quantified by qPCR targeting the 16S rRNA gene using primers Bac338F (5′-ACTCCTACGGGAGGCAGCAG-3′)^28^ and Bac534R (5′-ATTACCGCGGCTGCTGGC- 3′)^29^ on a QuantStudio™ 5 Real-Time PCR System with SYBR Green chemistry.

### 2.4 16S rRNA gene sequencing

Microbial community composition was characterized by 16S rRNA gene amplicon sequencing performed by Novogene (Netherlands) according to a previous study^30^.

### 2.5 Data analysis and visualization

#### 2.5.1 Calculation of gene abundances

Absolute abundances (AA) of ARGs and MGEs, expressed as gene copy numbers per gram of wet mass, were calculated by integrating relative gene abundances obtained from ResistoMap with absolute 16S rRNA gene copy numbers quantified by qPCR and DNA yields measured using a Qubit fluorometer. For substrate mixtures composed of multiple input materials, AA values were calculated as the mass-weighted sum of the AA of the individual substrates. Relative abundances (RA) of substrate mixtures were subsequently derived by normalizing the AA values to corresponding 16S rRNA gene abundances within the same mixtures.

#### 2.5.2 Multivariate and ordination analyses

Amplicon sequencing data of the 16S rRNA gene were processed using DADA2 (version 1.16) following established pipelines.^31^ Non-metric multidimensional scaling (NMDS) was used to visualise similarities among biogas digesters based on physicochemical characteristics and microbial community composition. NMDS of physicochemical parameters (pH, NH_3_–N, NH_4_^+^–N, TS, VS, OLR, HRT, and temperature) was performed using Euclidean distance, while NMDS of microbial community composition was conducted using Bray–Curtis dissimilarity. All ordinations were performed with two dimensions (k = 2), and random seeds were set to ensure reproducibility.

#### 2.5.3 Network analysis of host–resistance–mobility associations

Prior to network construction, a step-wise feature filtering strategy was applied to reduce sparsity and multiple-testing burden. ARGs and MGEs detected in fewer than 30% of samples were excluded (≥13/42). Microbial taxa were filtered by taxonomic resolution, with genera detected in fewer than 30% of samples and phyla detected in fewer than 10% of samples removed. To further improve statistical power, retained features were reduced to the most abundant sets, including the top 50 bacterial genera ranked by mean RA and the top 80 ARG subtypes ranked by total RA across all samples. Phylum-level taxa and MGEs were not further reduced by abundance, as their feature numbers were already limited following prevalence-based filtering. Relative abundance tables were centred log-ratio (CLR) transformed prior to correlation analysis, which is widely recognised as appropriate for compositional microbiome and gene-abundance data.^32^ Multiple testing was addressed using false discovery rate (FDR) correction, which provides a more stringent control of false positives than nominal p values in large-scale correlation-based network analyses.^33^

For stage-wise association dynamics, Spearman correlation analysis (|ρ| ≥ 0.6, FDR-adjusted p < 0.05) was applied to CLR-transformed data across all samples to identify significant genus– ARG, genus–MGE, and ARG–MGE associations. These associations were subsequently evaluated within individual process stages (substrate, digestate, and post-digestion storage) using a slightly relaxed yet conservative correlation threshold (|ρ| ≥ 0.5, FDR-adjusted p < 0.05), allowing assessment of the retention, loss, or emergence of correlations across the AD process without reconstructing networks from small stage-specific subsets. In addition, conventional tripartite correlation networks linking microbial taxa at the phylum level with ARGs and MGEs were constructed using Spearman correlation (|ρ| ≥ 0.6, FDR-adjusted p < 0.05) and visualized using Gephi (version 0.10.1).

#### 2.5.4 Statistical analysis of directional shifts in ARGs and MGEs

Changes in the RA of ARGs and MGEs were quantified for each biogas plant as Δ = (downstream process stage − upstream stage: Digestate – Substrate; Storage − Digestate). For each biogas plant, Δ values were first calculated at the subtype level. The classification of 323 ARG subtypes and 52 MGEs is summarised in Table S3. For class-level analyses, subtype-level Δ values were aggregated within each plant by calculating the median for each ARG class or MGE category. These class-level Δ values were then evaluated across plants using two-sided Wilcoxon signed-rank tests against zero. For overall ARG and MGE analyses, subtype-level Δ values were further aggregated within each plant by calculating the median across all subtypes prior to statistical testing. Missing Δ values, corresponding to resistance determinants not detected at a given process stage within a system, were excluded on a per-comparison basis. All analyses were performed in R. Given the observational nature of full-scale biogas plants and the absence of biological replicates, statistical analyses were restricted to evaluating directional trends in ARG and MGE abundances across digesters, rather than inferring absolute removal or enrichment.

## 3. Results

### 3.1 Abundance changes of ARGs and MGEs across anaerobic digestion systems

Across all substrate mixes, the absolute abundance (AA) of ARGs ranged from 3.44 × 10^8^ to 2.07 × 10^10^ gene copies per gram of wet mass, while MGEs ranged from 5.39 × 10^8^ to 1.03 × 10^10^ gene copies per gram of wet mass. No significant differences in the AA of ARGs or MGEs were observed among substrate mixes derived from different farming practices (Fig. 1A). In contrast, changes in AA throughout the anaerobic digestion (AD) system varied substantially among biogas plants, with both increases and decreases observed (Fig. 1B). During AD, decreases in AA were observed in plants H1, K2, and S, whereas increases occurred in plants L, V, and J (Table 1). During post-digestion storage, reductions in AA were detected in plants K1, K2, and N, while increases were observed in plants L, V, J, and H2 (Table 1).

**Table 1.** Percentage changes in absolute abundances of antibiotic resistance genes (ARGs) and mobile genetic elements (MGEs) during anaerobic digestion (Digester) and post-digestion storage (Storage), calculated relative to the corresponding substrates (Digester – Substrate; Storage – Substrate).

| No. | Digester | ARGs | MGEs | No. | Storage | ARGs | MGEs |
| --- | --- | --- | --- | --- | --- | --- | --- |
| 1 | H1_D1 | -51% | -72% | 1 | H2_S | 812% | 28% |
| 2 | H1_D2 | -81% | -83% | 2 | L_S | 2226% | 1401% |
| 3 | H2_D | 767% | 46% | 3 | V_S | 1024% | 84% |
| 4 | L_D | 1848% | 1033% | 4 | B_S | 614% | 147% |
| 5 | V_D1 | 1080% | 30% | 5 | K2_S1 | -82% | -91% |
| 6 | V_D2 | 2042% | 170% | 6 | K2_S2 | -90% | -95% |
| 7 | B_D | 396% | 227% | 7 | J_SS | 4092% | 1055% |
| 8 | K2_D | -13% | -26% | 8 | J_LS | 478% | 5% |
| 9 | J_D1 | 988% | 52% | 9 | K1_S | -64% | -89% |
| 10 | J_D2 | 1085% | 481% | 10 | N_S | -68% | -55% |
| 11 | J_D3 | 804% | 90% |  |  |  |  |
| 12 | K1_D | 117% | -28% |  |  |  |  |
| 13 | S_D | 8% | -51% |  |  |  |  |
| 14 | S_SD | -53% | -70% |  |  |  |  |
| 15 | N_D | -30% | 2% |  |  |  |  |

Absolute abundances refer to gene copy numbers per gram of wet mass.

**Fig. 1.**
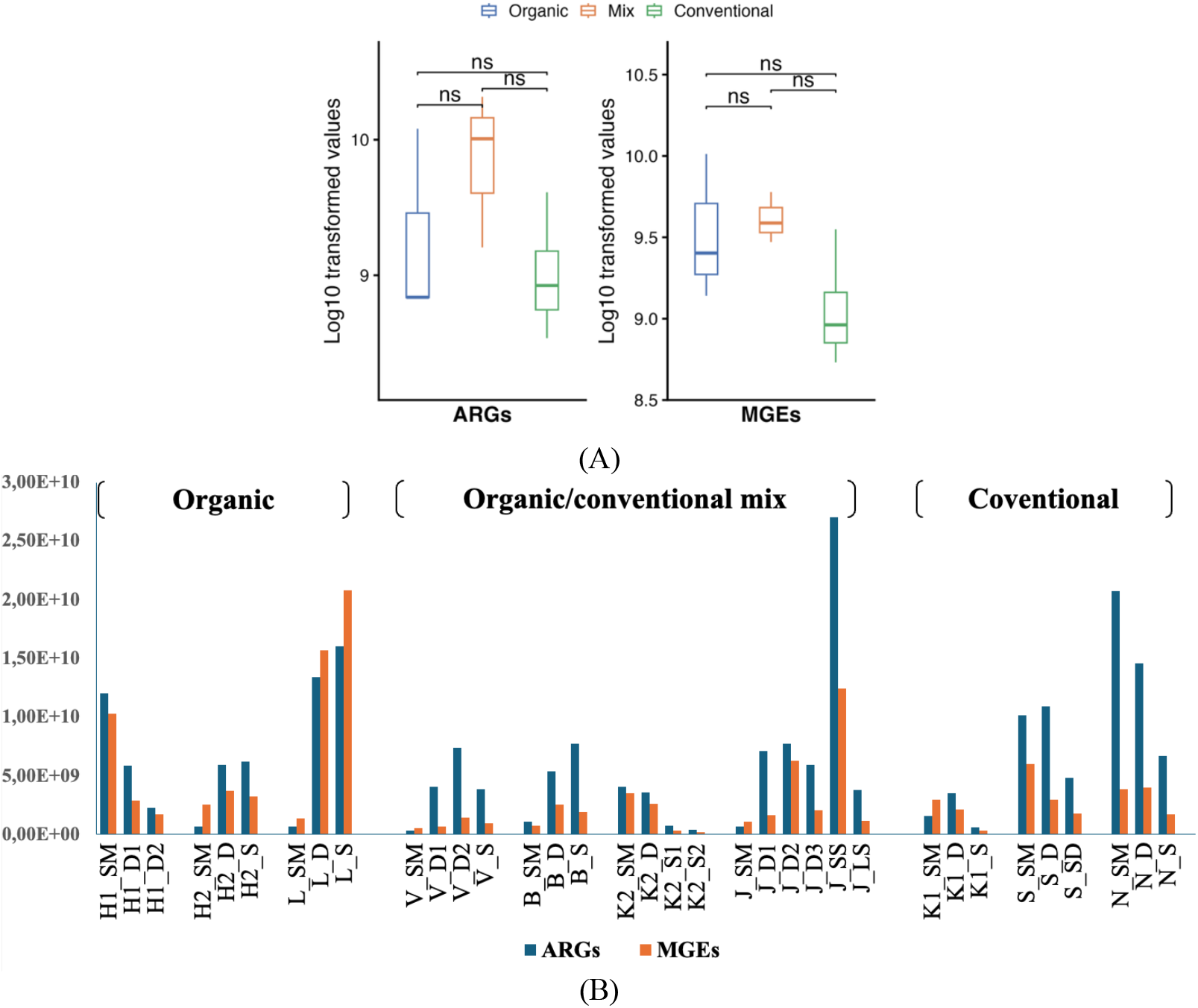
Absolute abundance of antibiotic resistance genes (ARGs) and mobile genetic elements (MGEs), expressed as gene copy numbers per gram of wet mass. (A) Abundance variation of ARGs and MGEs in substrates grouped by farming practices (organic, conventional, and mixed). Pairwise differences among farming practices were evaluated using two-sided Wilcoxon rank-sum tests. (B) Abundance dynamics across anaerobic digestion and post-digestion storage in ten full-scale biogas plants.

Changes in the relative abundance (RA) of ARG subtypes exhibited heterogeneous responses to AD, with both positive and negative shifts observed across antibiotic classes and digesters (Fig. 2A). Compared to substrate, none of the ARG classes showed a statistically significant directional shift in RA across the 15 digesters (Wilcoxon signed-rank test against zero, FDR-adjusted p > 0.05 for all classes), indicating that no specific ARG class was more likely to be enriched or reduced during AD. When RA changes across all ARG subtypes were aggregated, digesters exhibiting negative and positive Δ values generally corresponded to those showing decreasing and increasing trends in AA, respectively. However, when examined across all 15 digesters as a whole, the median change in RA did not deviate from zero (median Δ = 5.38 × 10^−6^; Wilcoxon signed-rank test against zero, FDR-adjusted p = 0.798), indicating the absence of a consistent directional shift in ARG RA during full-scale AD.

**Fig. 2.**
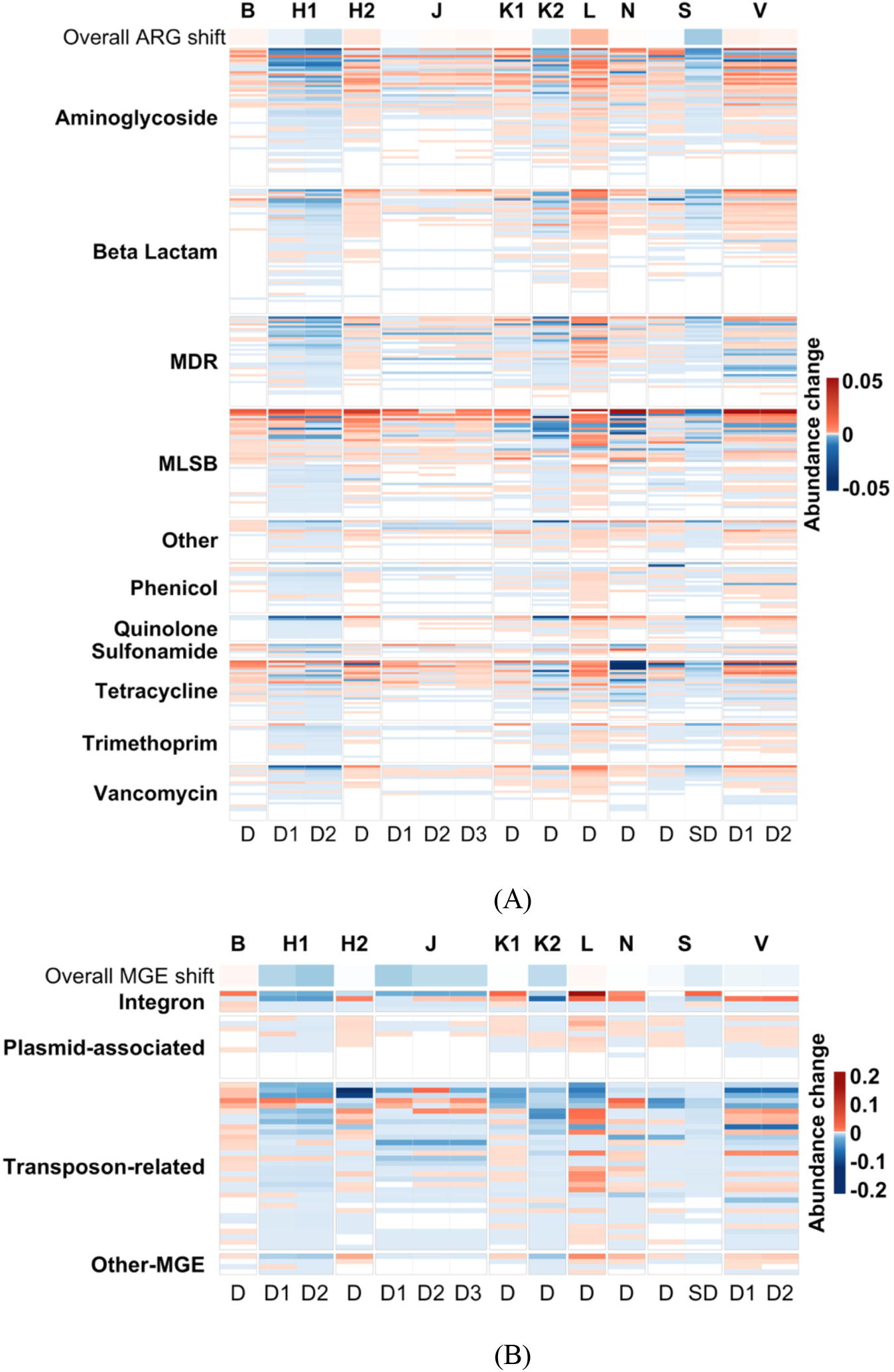
Fate of antibiotic resistance genes (ARGs) and mobile genetic elements (MGEs) during anaerobic digestion in ten full-scale biogas plants. Changes in relative abundance are expressed as the difference between digester and corresponding substrate samples (digester – substrate; for the digester S_SD, values are expressed as SD – D). Red and blue indicate increases and decreases in relative abundance, respectively. (A) ARG subtypes grouped by antibiotic classes. (B) MGEs grouped by functional categories. Columns represent individual digesters; D and SD denote digester and secondary digester, respectively. For both panels, the top bar summarizes the overall median change (Δ) in relative abundance for each digester, where blue and red indicate negative and positive deviations, respectively. The colour scale of the top bar is independent from that of the main heatmap and is used solely to indicate the direction of overall change.

In contrast to ARGs, distinct patterns were observed for RA changes of MGEs during AD (Fig. 2B). Transposon-related MGEs exhibited a statistically significant negative directional shift in RA across digesters (median Δ = −2.19 × 10^−4^, Wilcoxon signed-rank test against zero, FDR-adjusted p < 0.01). When normalised to baseline RA in substrate mixes, this shift corresponded to approximately 33% of the median baseline level (Table S4). No significant directional shifts were detected for integrons, plasmid-associated MGEs, or other MGEs, whose median Δ values remained close to zero and showed substantial variability among digesters. When RA changes across all MGEs were aggregated, despite variability among plants, a statistically significant negative directional shift was observed (median Δ = −1.19 × 10^−4^; Wilcoxon signed-rank test against zero, FDR-adjusted p < 0.01; Table S4), corresponding to approximately 25% of the median baseline abundance in substrate mixes (Table S4). Similar to the consistency between AA and RA changes observed for ARGs during AD, RA changes of MGEs were generally consistent with corresponding trends in AA. Exceptions were observed in plants J and V, where a negative shift in RA coincided with an increase in AA. This discrepancy was likely driven by a substantial increase (6–10-fold) in absolute 16S rRNA gene abundance in digestates relative to substrates in these plants. Consequently, despite a negative shift in RA, AA increased due to the overall proliferation of the microbial community.

Regarding post-digestion storage, no consistent directional shifts in RA of ARGs or MGEs were observed across the ten storage treatments (Fig. S1). Median Δ values for ARG classes and MGE categories remained close to zero, indicating that storage did not introduce systematic enrichment or depletion beyond the post-digestion state. Notably, in plant J, where liquid–solid separation of digestate was applied, the RA of ARGs and MGEs was consistently higher in the solid fraction than in the corresponding liquid fraction derived from the same digestate (Fig. S1). Consistent with this pattern, AA of ARGs and MGEs was also substantially higher in the solid fraction than in the liquid fraction (Table 1). Together, these results indicate a preferential association of ARGs and MGEs with the solid fraction of digestate.

### 3.2 Effect of physicochemical parameters on ARG and MGE dynamics during anaerobic digestion

Physicochemical parameters, including TS, VS, pH, calculated free ammonia (NH_3_), ammonium nitrogen (NH_4_^+^-N), VFAs, HRT, temperature and OLR, were evaluated to explore their potential associations with changes in the RA of ARGs and MGEs during AD. For ARGs, samples exhibiting overall decreasing trends in RA tended to cluster in the direction of higher pH and ammonia-related parameters, as reflected by the positioning of samples such as K2_D, H1_D1, and H1_D2 along the direction of the pH- and ammonia-related vectors (Fig. 3A). This pattern suggests a potential association between elevated pH and ammonia levels and reductions in overall ARG RA. An exception was observed for S_SD, a secondary digester, which deviated from this general trend. For MGEs, most digesters displayed overall decreasing trends in RA, with only B_D and L_D showing increasing median RA values (Fig. 2B). These samples were positioned opposite to the pH and ammonia-related vectors, consistent with the pattern observed for ARGs (Fig. 3B). Overall, the ordination results indicate that pH and NH_3_ are associated with decreasing trends in the RA of ARGs and MGEs across the investigated full-scale anaerobic digesters.

**Fig. 3.**
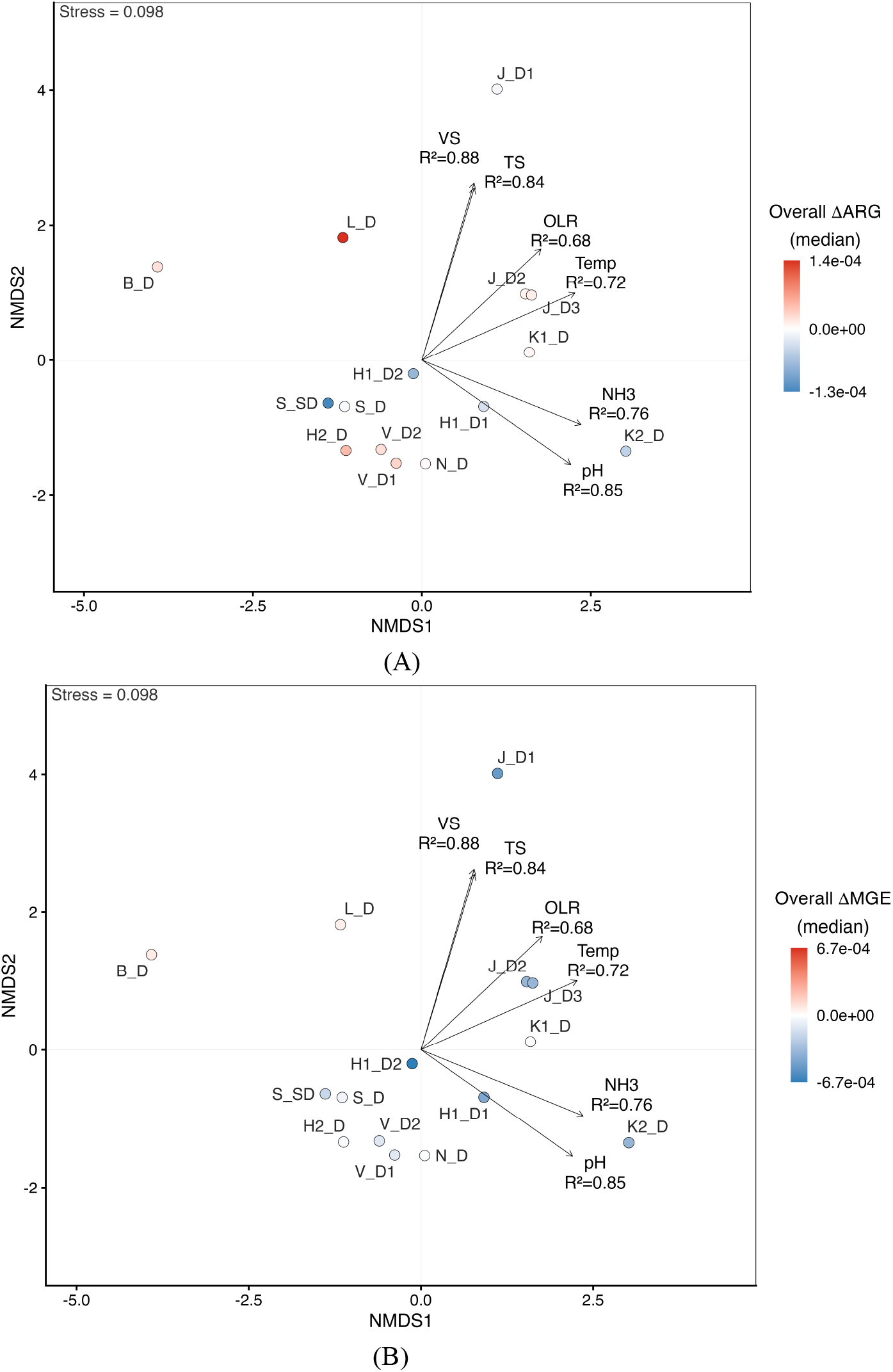
Non-metric multidimensional scaling (NMDS) ordination of full-scale biogas digesters based on physicochemical parameters, with samples coloured by overall median shifts. (A) Samples coloured by the overall median change in antibiotic resistance genes (ΔARG). (B) Samples coloured by the overall median change in mobile genetic elements (ΔMGE). Colour gradients range from blue (negative change) to red (positive change), with white indicating no change. Arrows indicate relationships between physicochemical parameters and sample ordination. Only variables showing significant correlations (p ≤ 0.05) are shown.

### 3.3 Network patterns of host–ARG–MGE associations

#### 3.3.1 Overview of the host–ARG–MGE network structure

A tripartite network integrating samples from substrates, digestates and post-digestion storages was constructed to capture associations among ARGs, MGEs and microbial hosts across full-scale biogas systems (Fig. 4). Based on robust positive correlations ((|ρ| ≥ 0.6, FDR-adjusted p < 0.05), the network comprised 67 ARG nodes, 30 MGE nodes, and 18 microbial phylum nodes, connected by a total of 204 edges. Distinct levels of connectivity were observed among the three network components, with MGEs exhibiting the highest number of associations, followed by ARGs, while microbial phyla showed the lowest connectivity. Among MGEs, several high-degree nodes were identified, including orf37-IS26 (n = 21), IS21–ISAs29 (n = 15), and ISCR1 (n = 12). Similarly, several ARGs displayed relatively high connectivity, such as blaOXY (n = 12), APH(3’)-Ib (n = 11), and rmtB (n = 7). In contrast, microbial phylum nodes exhibited substantially fewer associations, with Spirochaetota (n = 8), Abditibacteriota (n = 7), and Desulfobacterota (n = 3) representing the most connected phyla. Overall, the tripartite network revealed that ARG–MGE associations formed the dominant structural backbone, whereas microbial phyla occupied more peripheral positions, reflecting limited but specific associations.

**Fig. 4.**
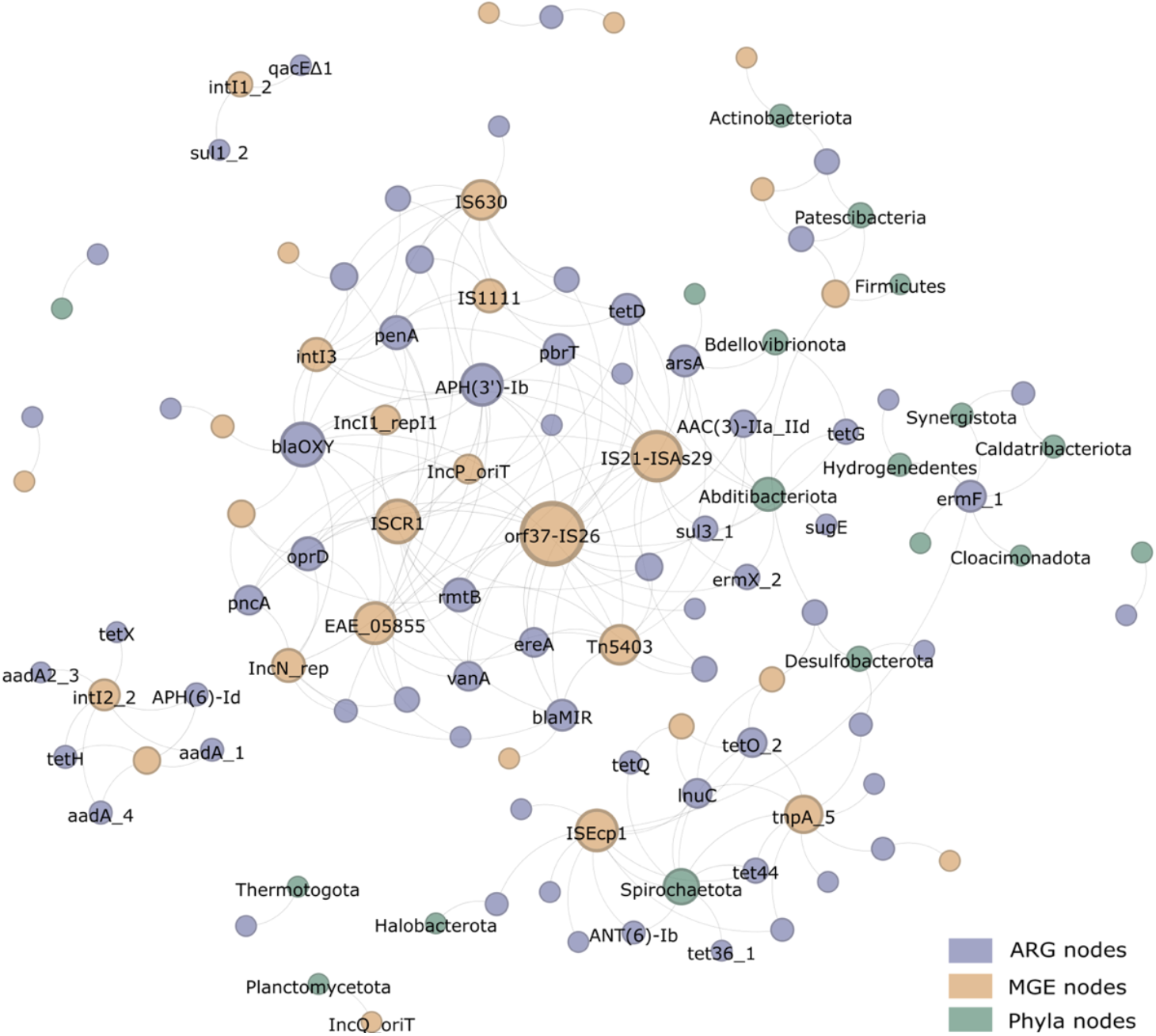
Tripartite network integrating antibiotic resistance genes (ARGs), mobile genetic elements (MGEs), and microbial phyla based on robust positive correlations (ρ > 0.6, false discovery rate (FDR)-adjusted p < 0.05). Node colours indicate ARGs, MGEs, and microbial phyla. Node size is proportional to node degree. Node labels are shown only for highly connected nodes or those discussed in the main text to improve visual clarity.

#### 3.3.2 Stage-wise dynamics of host–ARG–MGE associations

To characterise the dynamics of associations among hosts, ARGs, and MGEs across process stages, a global bipartite network was inferred from 42 samples, encompassing 17 substrates, 15 digestates, and 10 post-digestion storage samples. The resulting associations were subsequently evaluated within individual process stages (substrate, digestate, and post-digestion storage) to determine whether they were retained, lost, or newly supported across stages (Fig. 5). Microbial associations with ARGs and MGEs were assessed at the genus level to provide sufficient host resolution.

**Fig. 5.**
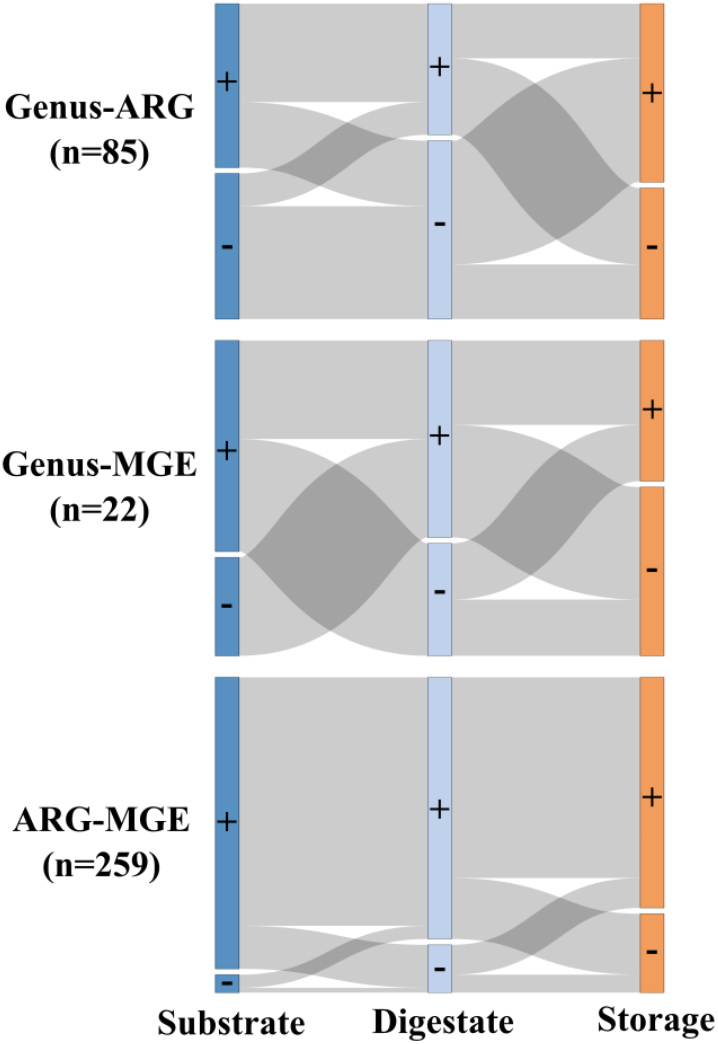
Bipartite Sankey diagrams illustrating stage-wise changes in Genus–ARG, Genus–MGE, and ARG–MGE associations across substrate, digestate, and storage stages. Bipartite associations were first inferred from the full dataset using all samples, and the resulting association pool was subsequently evaluated for stage-specific support. Symbols “+” and “–” indicate whether an association was supported or not supported at a given stage. Flows represent the retention, loss, or emergence of associations between consecutive stages, reflecting stage-wise restructuring of association pools rather than persistence of individual links. Numbers in parentheses indicate the total number of associations included in each bipartite network.

For genus–ARG associations, stage-wise analysis revealed pronounced restructuring across the AD system (Fig. 5). From substrate to digestate, a substantial proportion of identified genus– ARG associations was lost, accompanied by the emergence of a comparable number of new associations, reflecting marked shifts in ARG host affiliations during the digestion process. This dynamic pattern became more pronounced during the transition from digestate to post-digestion storage, where newly formed genus–ARG associations exceeded retained ones and additional losses were observed. Together, these changes demonstrate continued reorganisation of ARG–host relationships during post-digestion storage rather than stabilisation after digestion. Topological analysis of the genus–ARG network showed a relatively even distribution of node degrees across bacterial genera, with no single genus acting as a dominant hub taxon (Table S5). Instead, ARG associations were distributed among multiple taxa, suggesting frequent reassortment of ARG–host linkages across successive process stages.

Compared with genus–ARG networks, genus–MGE associations were fewer in number and exhibited intermediate levels of stability across stages (Fig. 5). From substrate to digestate, multiple host–MGE associations were lost while new associations concurrently emerged. During the transition from digestate to storage, both retained and newly formed genus–MGE associations were observed, accompanied by fewer losses compared to substrate-digestate transition. The continued appearance of new host–MGE associations indicates ongoing redistribution of MGEs among microbial taxa beyond the digestion stage. Network topology analysis further showed that genus–MGE associations were markedly sparse, with most genera and MGEs exhibiting low connectivity and no pronounced hub structure (Table S4). This degree pattern reflects generally limited and transient host–MGE associations at the genus level.

In contrast, ARG–MGE associations exhibited a markedly different stage-wise pattern compared with host-associated networks (Fig. 5). From substrate to digestate, the majority of ARG–MGE links were retained, with relatively few losses or newly formed associations, indicating strong conservation of resistance–mobility coupling during anaerobic digestion. This high degree of stability persisted during the transition from digestate to storage, where retained ARG–MGE associations dominated and only limited gains or losses were observed.

In contrast to the dynamic restructuring of genus–ARG and genus–MGE associations, ARG– MGE coupling remained largely conserved across stages despite changes in microbial host composition. Topological analysis of the ARG–MGE network indicated that connectivity was driven by a small number of highly connected MGEs (e.g., IS26-, ISEcp-, and IS21-associated elements; Table S5). These MGEs were repeatedly associated with multiple different ARGs across stages, whereas no individual ARG subtype exhibited comparable connectivity. Together, this indicates that ARG–MGE associations across stages are primarily structured by MGEs rather than by ARG-specific connectivity.

Together, these network analyses indicate pronounced restructuring of host–ARG and host– MGE associations across process stages, while ARG–MGE coupling remains comparatively stable. To further characterise these network-derived associations, the RA patterns of the associated bacterial genera were examined.

### 3.4 Abundance patterns of ARG- and MGE-associated bacterial genera

To place these network-associated bacterial genera in an abundance context, the RA of ARG- and MGE-associated genera was examined across samples and process stages (Fig. 6). As shown in Fig. 6, the majority of ARG- and MGE-associated bacterial genera occurred at appreciable RA, indicating that these associations are not driven solely by rare taxa. Overall, the number of ARG-associated genera was higher than that of MGE-associated genera. In total, these genera belonged to six bacterial phyla, with Firmicutes representing the dominant group, accounting for 18 out of 30 genera. The RA of ARG- and MGE-associated genera varied across samples and process stages (Fig. 6). Although no consistent RA trends for these genera were observed across process stages, many genera exhibited relatively limited RA fluctuations, particularly those with high network degrees identified in the network analysis (Table S5). For example, the RA of *Acholeplasma* (n=13) and *Syntrophomonas* (n=8) remained comparatively stable across stages. In contrast, genera with lower network degrees showed more pronounced RA variability across process stages, including genera such as *Pantoea* and *W5*.

**Fig. 6.**
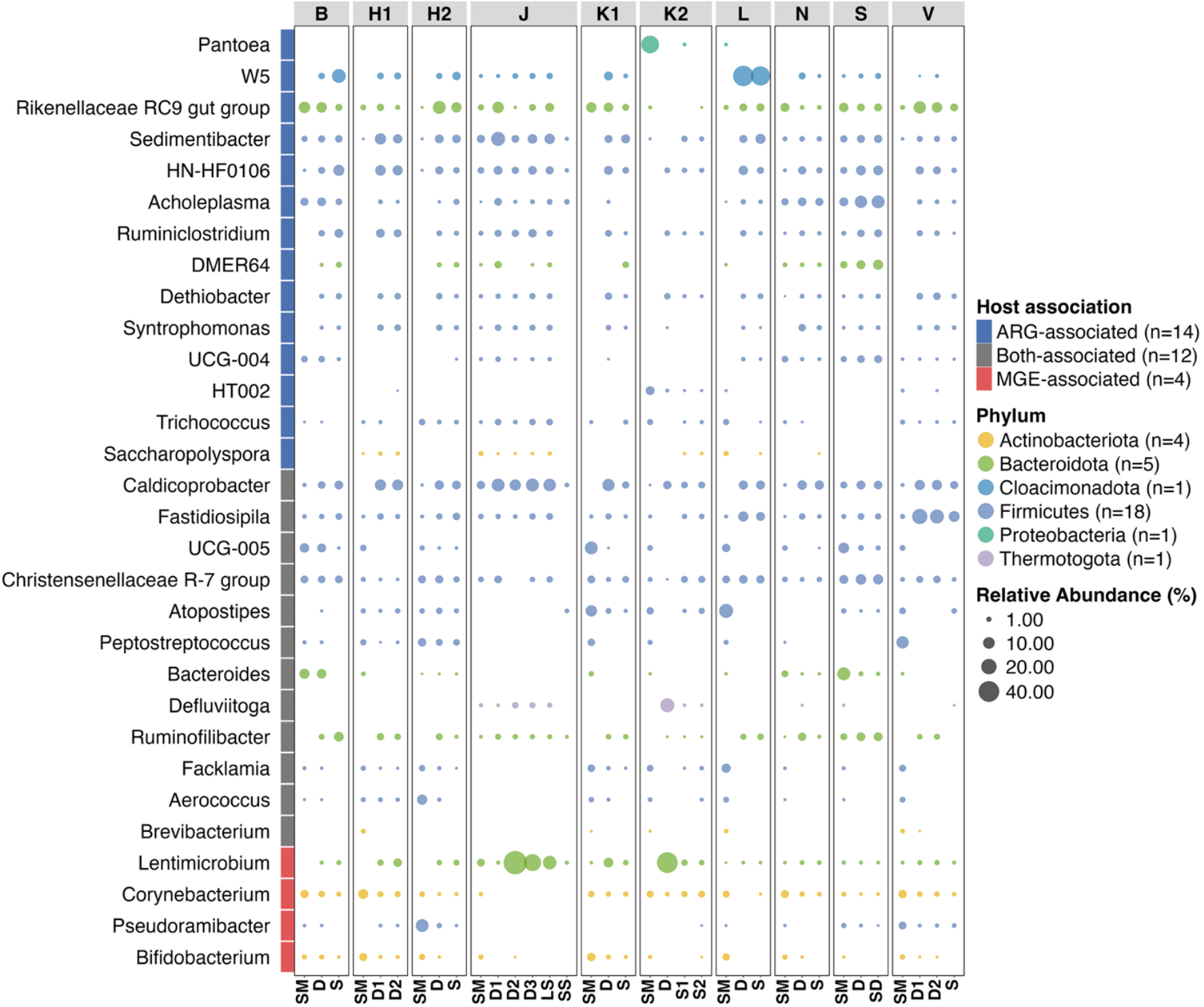
Distribution of bacterial genera associated with antibiotic resistance genes (ARGs) or mobile genetic elements (MGEs) across full-scale biogas digesters. Genera are classified as ARG-associated, MGE-associated, or associated with both, as indicated by the annotation strip on the left; numbers in parentheses denote the total number of genera in each association category. Bubble size represents genus-level relative abundance, while colors denote bacterial phyla; numbers in parentheses indicate the total number of genera detected within each phylum. For visual clarity, data points with relative abundance <0.1% are omitted from bubble display.

## 4 Discussions

### 4.1 Abundance level of ARGs and MGEs in agricultural substrates under low-antibiotic-use conditions

Nordic countries have some of the strictest regulations on antibiotic use in agriculture worldwide, and Sweden was the first country to ban antibiotic growth promoters in animal feed as early as 1986.^34^ Under such stringent regulatory conditions, the similar absolute abundance (AA) levels of ARGs and MGEs observed in substrates derived from different farming practices—including organic, conventional, and mixed conventional–organic systems— suggest that differences in antibiotic resistance loading associated with substrate origin are limited. Although the AA of selected ARGs has been reported in previous anaerobic digestion studies,^35–37^ comprehensive assessments of the overall AA of ARGs across a broad spectrum of resistance genes remain scarce, particularly in full-scale, agriculture-based biogas systems. As a result, the AA levels of ARGs and MGEs in substrates from the ten farm-scale biogas plants investigated here cannot be directly compared with existing studies. Nevertheless, the AA data generated in this study provide valuable baseline information on resistome-wide ARG and MGE loads in agricultural substrates entering full-scale AD systems under low-antibiotic-use conditions.

### 4.2 Effect of physicochemical parameters on abundance changes in ARGs and MGEs during anaerobic digestion

Across the investigated farm-scale AD systems, changes in AA and RA of ARGs and MGEs generally followed consistent directional patterns, with exceptions only in a limited number of plants where MGE exhibited divergent trends between AA and RA. At the system level, neither AA nor RA of ARGs showed a uniform directional trend across the 15 investigated digesters, with both increases and decreases observed depending on site-specific conditions. This heterogeneity indicates that AD cannot be assumed to universally reduce ARG abundance under real-world operational settings. Instead, the fate of ARGs appears to be highly context-dependent, shaped by plant-specific configurations and operating conditions as discussed below.

In the present study, samples exhibiting overall decreasing trends in ARG and MGE abundances were generally positioned along gradients of higher pH and NH_3_ in the ordination analysis, indicating a potential association between these physicochemical conditions and the attenuation of resistance and mobility determinants (Fig. 3). Notably, pH and NH_3_ concentrations are intrinsically linked in AD systems, as increases in pH are associated with changes in ammonia speciation toward a higher proportion of NH_3_. Previous studies have demonstrated that the influence of AD physicochemical parameters on ARG and MGE dynamics is predominantly mediated through changes in the microbial community, rather than through direct effects on genes themselves.^16,21^ In this context, NH_3_ has been reported to exert strong selective pressure on microbial community composition.^38^ Elevated NH_3_ concentrations have been shown to reduce microbial diversity during AD,^39,40^ and lower microbial diversity has, in turn, been associated with greater reductions in ARGs, likely reflecting a reduced availability of potential microbial hosts.^15,16^ More direct evidence for the role of NH_3_ in ARG attenuation has also been provided by an experimental study were free ammonia pretreatment (420 mg NH_3_–N L^−1^ for 24 h) significantly enhanced the reduction in absolute abundance of multiple ARGs, including AAC(6′)-Ib-cr, blaTEM, sul2, tetA, tetB, and tetX, during subsequent anaerobic digestion^41^. Consistent with this finding, our previous study also observed enhanced ARG reduction under higher NH_3_ levels in agricultural biogas plants treating plant-based substrates.^9^

However, elevated pH and NH_3_ levels alone were not determinative of ARG and MGE dynamics. In the present study, for example, digesters H1_D2 and V_D2 exhibited divergent fates of ARGs and MGEs, with H1_D2 showing an overall reducing pattern and V_D2 exhibiting an enriching pattern (Table 1), despite comparable pH and NH_3_ concentrations (Table S2). This indicates that combined pH–NH_3_ conditions were insufficient to fully explain resistance gene dynamics. This divergence of gene fates likely reflects the influence of additional physicochemical parameters acting in concert with pH and NH_3_. Among these factors, temperature is one of the most extensively studied and widely regarded as a critical parameter shaping microbial community structure in AD systems.^10,42^ Numerous studies have reported that thermophilic AD generally achieves greater reductions in ARGs than mesophilic operation.^11,16,17^ For instance, Diehl and LaPara^43^ demonstrated that increasing digestion temperature from 22 °C to 55 °C significantly enhanced the removal of tetracycline resistance genes and the class I integron integrase gene *intI1*, highlighting the sensitivity of both ARGs and integron-associated elements to thermal conditions. More recently, Zou et al.^14^ showed that thermophilic digestion (55 °C) promoted the reduction of extracellular ARGs and *intI1* compared with mesophilic operation (35 °C), suggesting that elevated temperature may facilitate the degradation of both intracellular and extracellular genetic material. In this context, the contrasting ARG and MGE trajectories observed in digesters H1_D2 and V_D2 may be partly attributable to differences in operating temperature, with H1_D2 operating at 41 °C compared with 38 °C in V_D2. In addition to temperature, other physicochemical parameters may also have contributed to the divergent ARG and MGE fates observed in H1_D2 and V_D2, including differences in OLR, VS, and TS, as suggested by the ordination analysis (Fig. 3). However, existing studies provide limited evidence for consistent or direct associations between these parameters and ARG or MGE reduction under typical wet AD conditions. Notably, enhanced ARG and MGE reduction associated with higher TS has primarily been reported for high-solids or dry anaerobic digestion systems (TS > 15%), which have been shown to outperform wet-type systems (TS < 8%) in terms of ARG attenuation.^20,21^ In the present study, TS values in both H1_D2 and V_D2 remained below 8%, suggesting that differences in TS within this low range are unlikely to represent a dominant driver of the contrasting resistance gene dynamics observed between these digesters, despite the relatively higher TS in H1_D2.

Collectively, among the physicochemical parameters examined across the 15 digesters, elevated pH and NH_3_ were the parameters most consistently associated with effective ARG and MGE reduction, likely through their influence on microbial community structure. Nevertheless, the combined pH–NH_3_ condition alone was not determinative of ARG and MGE dynamics, and the observed variability among digesters indicates that multiple physicochemical parameters act in concert to shape ARG and MGE fate during full-scale AD.

### 4.3 Effect of system configurations on abundance changes in ARGs and MGEs

#### 4.3.1 Two-stage digestion

Beyond physicochemical parameters, process configuration has also been shown to play an important role in shaping the dynamics of ARGs and MGEs during anaerobic digestion. Two-phase digestion systems have repeatedly been reported to achieve greater reductions in ARGs and intI1 than single-stage systems, with the acidogenic phase often identified as a critical window for resistance attenuation, followed by more variable responses during the methanogenic stage.^19,44^ Evidence from full-scale systems further supports this pattern. For example, Jang et al.^18^ reported significant reductions of ARGs and intI1 in two-phase thermophilic digesters, with several resistance genes becoming undetectable in the final effluent. It should be noted that the secondary digesters examined in the present study does not represent a classical acidogenic–methanogenic two-phase configuration. In plant S, the primary digester operated at near-neutral pH (7.67) with a relatively long hydraulic retention time (29 d), during which the main anaerobic digestion processes were already completed. The secondary digester (pH 7.48; HRT 4 d) therefore functioned primarily as a post-digestion polishing unit by extending the overall residence time, rather than introducing a distinct metabolic phase. Despite the different configuration, the two-stage system in plant S still exhibited pronounced reductions in ARGs and MGEs. One possible explanation is the comparably more oligotrophic conditions in the secondary reactor, which has been associated with reduced microbial diversity and enhanced ARG removal.^45,46^ This interpretation is supported by the comparison of microbial community structures between the primary and secondary digesters (Fig. S2), where both the relative abundance and diversity of genera appeared to be lower in the secondary digester compared with the primary digester. In addition, prolonged residence time may promote antibiotic biodegradation,^47^ potentially resulting in lower residual antibiotic concentrations in the secondary digester and, consequently, reduced selective pressure for antibiotic resistance. Overall, although the two-stage configuration in plant S differs from a classical acidogenic–methanogenic system, it nevertheless exhibited a comparably effective reductions of ARGs and MGEs. This finding may suggest that extending residence time through multi-stage configurations may also lead to enhanced ARG and MGE reduction, similar to outcomes reported for classical two-phase digestion. In this context, two-stage operation can represent a potentially effective approach for reducing ARGs and MGEs.

#### 4.3.2 Post-digestion storage

In full-scale biogas systems, post-digestion storage is routinely implemented to balance temporal mismatches between digestate production and agricultural application, to stabilise digestate properties, and to facilitate subsequent handling practices such as solid–liquid separation.^48,49^ Beyond these operational functions, post-digestion storage has also been recognised as a stage at which further changes in microbial activity and associated genetic elements may occur. ^50,51^

Post-digestion storage exhibited heterogeneous effects on ARG and MGE abundances in the present study, with both increases and decreases observed across storage treatments in terms of both absolute abundance (AA) and relative abundance (RA). Such variability is consistent with previous studies, which have shown that ARG dynamics during storage are highly gene- and subtype-specific rather than uniform. For example, Zhang et al.^50^ reported that during 30 days of digestate storage, total ARG RA decreased overall, while several ARG subtypes—including tetM, tetX, tetQ, tetS, ermF, and sul2—persisted or increased. Similarly, Pu et al.^51^ observed pronounced enrichment of ARGs in RA during 28 days of aerobic storage of biogas residues. Together, these findings indicate that individual resistance genes may exhibit divergent responses during storage, even when overall abundance trends appear limited or inconsistent.

Beyond gene-level heterogeneity, post-digestion storage has also been reported to influence the partitioning of ARGs between solid and liquid fractions. Zhang et al.^49^ showed that following AD of dairy manure, ARG AA decreased in the liquid fraction but was concurrently enriched in the solid fraction, with higher total solids content associated with increased ARG concentrations in solids. This pattern is consistent with our observations from solid–liquid separated storage, where both AA and RA of ARGs were consistently higher in the solid fraction than in the corresponding liquid fraction (Table 1 and Figure S2). Collectively, these results suggest that post-digestion storage represents a complex and heterogeneous stage with respect to ARG and MGE dynamics, characterised by gene-specific responses and phase-dependent distributions rather than uniform enrichment or reduction.

### 4.4 Network analysis of host–resistance–mobility associations

Reliable inference of microbiome association networks critically depends on adequate sample size and appropriate statistical control, particularly in high-dimensional datasets involving microbial taxa, ARGs and MGEs. Recent studies have cautioned that networks inferred from fewer than 20 samples per group are prone to instability and spurious associations.^52,53^ To address these constraints, the present study first constructed a global association network using all 42 samples thereby providing sufficient statistical power to identify robust associations across the AD system. Rather than reconstructing separate networks from small stage-specific subsets, stage-wise support of global associations was subsequently evaluated within individual process stages. Together with the application of CLR transformation and FDR– adjusted p-values,^32,33^ these methodological choices provide a stable and interpretable framework for inferring host–ARG–MGE associations in this study.

The conventional tripartite network highlighted a pronounced structural imbalance among hosts, ARGs, and MGEs (Fig. 4). ARG–MGE associations formed the dominant backbone of the network, whereas microbial hosts occupied more peripheral positions with comparatively fewer connections. This pattern is consistent with several previous network-based studies reporting limited host-ARG associations.^11,19,54^ In contrast, some studies have reported more extensive associations between microbial phyla and ARGs or MGEs.^21,24^ Such discrepancies may be attributable to methodological differences, including limited sample sizes (e.g., n = 6)^21^ or the use of less stringent correlation thresholds^24^, both of which can increase the likelihood of detecting spurious associations.

Network analyses in previous AD studies are commonly derived from aggregated samples and are often used to infer potential microbial hosts of ARGs or MGEs, thereby linking changes in ARG and MGE abundances to shifts in host community composition along the AD process. For example, enhanced ARG reduction has been attributed to the elimination of putative host bacteria during digestion.^55^ However, this interpretation is challenged by the dynamic host– ARG–MGE associations observed in the present study. Although ARG–MGE linkages remained comparatively stable across process stages, their microbial hosts shifted substantially from substrate to digestate and further to post-digestion storage. Consequently, association patterns inferred from aggregated samples represent static snapshots that may not be consistently supported at individual process stages. These findings suggest that network-based interpretations derived from aggregated datasets should be applied with caution when used to explain stage-specific ARG and MGE dynamics in AD systems.

An interesting observation emerged when host–ARG–MGE associations were examined at different taxonomic resolutions. Specifically, the identity of the most strongly associated phyla differed depending on whether hosts were defined at the genus or phylum level. At the genus level, the majority of highly associated taxa belonged to Firmicutes, followed by Actinobacteriota, which is consistent with findings reported in many previous studies.^24,54,55^ In contrast, when associations were analysed directly at the phylum level, Spirochaetota and Abditibacteriota emerged as the most highly associated phyla. In the present study, Spirochaetota exhibited associations with multiple ARGs, including tet44, tet36, tetQ, lnuC, and ANT(6)-Ib, as well as with MGEs such as ISEcp1 and tnpA (Fig. 4). *Treponema*, a representative belonging to Spirochaetota was also observed at relatively high abundance (Fig. S2). Consistent with previous reports, *Treponema* has frequently been associated with ARGs, particularly tetracycline resistance genes, in AD and related environments.^46,49,56^ By contrast, phylum Abditibacteriota has rarely been reported in AD systems, likely due to its generally low relative abundance (<1%). Nevertheless, it exhibited associations with multiple ARGs spanning different antibiotic classes, including tetD, tetG, sul3, ermX, sugE, arsA, and AAC(3)-IIa_IId, although no associations with MGEs were detected (Fig. 4). Despite its low abundance, Abditibacteriota has previously been reported to display high phenotypic resistance to a broad range of antibiotics.^57^ Overall, the inconsistency in top associated phyla across taxonomic levels likely reflects the fact that phylum-level taxa encompass diverse genera with distinct resistance-related roles. As a result, genus-specific host–ARG–MGE associations may be masked or diluted when analyses are conducted at broader taxonomic resolutions.

To our knowledge, this study represents the first application of a stage-resolved network framework to investigate ARG–MGE–host associations across the full AD and post-digestion storage process, moving beyond static correlation analyses based on aggregated samples. Using this approach, we show that while ARG–MGE associations remain largely conserved across process stages, microbial hosts associated with these resistance determinants undergo continuous reshuffling throughout the AD system. These findings suggest that ARG persistence in full-scale biogas plants is driven less by stable host lineages and more by the redistribution of ARG–MGE units among different microbial taxa. On this basis, we propose a conceptual framework in which anaerobic digestion functions primarily as a process of host reshuffling rather than complete resistance elimination (Fig. 7). This framework provides a mechanistic explanation for why abundance-based assessments alone often fail to resolve the apparent stability of the resistome across AD and post-digestion storage. Consistent with this interpretation, the absolute abundances of ARGs and MGEs in this study remained within a similar order of magnitude (10^8^–10^10^ gene copies per gram wet mass) from substrates through digestates and subsequent storage, despite pronounced changes in microbial community composition and host associations.

**Fig. 7.**
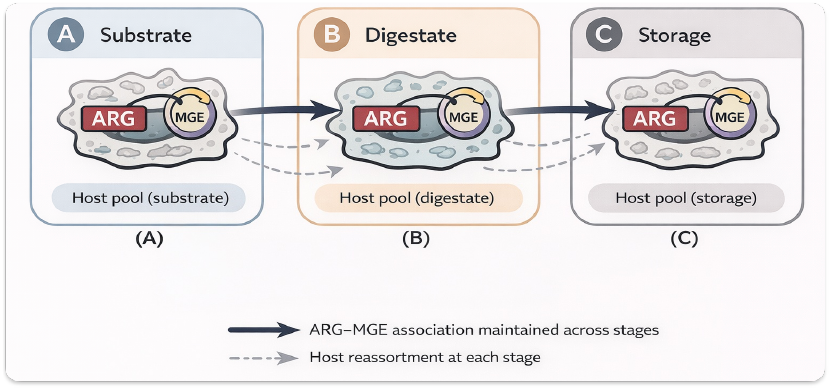
Conceptual framework illustrating the relationships among antibiotic resistance genes (ARGs), mobile genetic elements (MGEs), and microbial hosts across anaerobic digestion and storage stages. The scheme depicts ARG–MGE associations alongside stage-specific reassortment of microbial host pools.

### 4.5 Environmental implications

This study demonstrates that digestate storage should not be regarded as a passive holding phase prior to land application. Instead, storage represents a biologically active stage during which ARGs and MGEs may continue to be redistributed within microbial communities, reshaping host–gene associations beyond the digestion endpoint. Such dynamics imply that resistance determinants introduced into agricultural soils may reflect ongoing post-digestion processes rather than solely conditions at digester exit. At the same time, these effects should be interpreted in the context of the substantial initial ARG and MGE loads present in raw substrates. Against this large baseline, the moderate reductions or enrichments observed during anaerobic digestion and storage are unlikely to substantially alter absolute resistance inputs to the environment. Therefore, digestate remains an environmentally preferable alternative to untreated substrates such as raw manure, owing to its improved nutrient bioavailability, higher nutrient use efficiency, and comparable ARG and MGE levels relative to raw substrates. Overall, AD and subsequent storage do not eliminate resistance determinants but modify their ecological context prior to land application. Environmental risk assessments should therefore move beyond abundance-based metrics alone and incorporate post-digestion dynamics alongside agronomic benefits when evaluating digestate use in sustainable agricultural systems.

## 5 Conclusions

This study provides a stage-resolved assessment of ARG and MGE dynamics in full-scale AD systems. Across ten farm-scale biogas plants, ARGs and MGEs exhibited heterogeneous abundance changes during digestion and post-digestion storage. Elevated pH and NH_3_ conditions were associated with enhanced ARG and MGE reduction but were not determinative. Network analyses showed that ARG–MGE associations constitute a stable structural backbone across process stages, whereas microbial hosts are continuously reshuffled. Collectively, these findings demonstrate that resistance dynamics in AD systems cannot be adequately captured by static analyses. Stage-resolved, network-informed approaches therefore provide a more informative framework for evaluating resistance persistence and environmental risks associated with digestate application, and for informing sustainable biogas and nutrient recycling practices.

## Supporting information

All supplementary materials

