## Supplementary material for "Stage-resolved dynamics of antibiotic resistance genes and mobile genetic elements in full-scale anaerobic digestion systems": All supplementary materials

Table S1. Overview of samples collected from ten full-scale biogas plants in Sweden, including substrates and sampling points across the anaerobic digestion process.

| No. | Biogas plants | Substrates | | | | | | | Digestates | | | | | | | |
|---|---|---|---|---|---|---|---|---|---|---|---|---|---|---|---|---|
| | | Cow manure | Pig manure | Chicken manure | Deep bedding | Food waste | Slaughter-house waste | Plant materials | Primary digester1 | Primary digester2 | Primary digester3 | Secondary digester | Liquid separation | Solid separation | Storage1 | Storage2 |
| 1 | H1 | X | | X | X | | | X | X | X | | | | | | |
| 2 | H2 | X | | | | | | | X | | | | | | X | |
| 3 | L | O | O | | | | | X | X | | | | | | X | |
| 4 | V | O | O | O | | O | O | | X | X | | | | | X | |
| 5 | K2 | O | O | | △ | △ | | X | X | | | | | | X | X |
| 6 | B | O | O | | | | | | X | | | | | | X | |
| 7 | J | | | | | | | X | X | X | X | | X | X | | |
| 8 | K1 | X | | | | | | | X | | | | | | X | |
| 9 | S | O | | | O | | | | X | | | X | | | | |
| 10 | N | | O | | O | | | | X | | | | | | X | |

"X" indicates individual samples collected from a single substrate or process unit. "O" and "△" indicate samples obtained from premixed materials prior to sampling (i.e., composite samples representing multiple substrate inputs). Plant numbers highlighted in green (plants 1–3) represent organic farming systems, those in yellow (plants 4–7) represent mixed conventional–organic systems, and those in red (plants 8–10) represent conventional farming system.

Table S2. Operational parameters and sample characteristics of the ten full-scale biogas plants included in this study.

| Biogas plants | Samples | OLR[a], kg (VS[b])•$m^{-3}$•$d^{-1}$ | TS[c], % | VS, % of TS | pH | $NH_4^+$-N, mg•$L^{-1}$ | $NH_3$[d], mg•$L^{-1}$ | VFA[e], g•$L^{-1}$ | HRT[f], d | Temp, °C |
|---|---|---|---|---|---|---|---|---|---|---|
| H1 | Cow manure | 2.95 | 11.21±1.26 | 58.26±5.08 | 7.01 | 1142 | - | 7.54 | - | - |
| | Deep bedding | | 23.72±2.45 | 63.98±2.23 | - | - | - | - | - | - |
| | Oat water | | 15.70±1.62 | 69.66±2.04 | 4.44 | 115 | - | 10.12 | - | - |
| | Chicken manure | | 43.20±0.48 | 76.44±1.47 | - | - | - | - | - | - |
| | Digestate 1 | | 6.01±0.43 | 41.80±1.14 | 7.85 | 2800 | 294 | 0.95 | 28 | 41 |
| | Digestate 2 | | 5.46±0.21 | 38.98±1.14 | 7.45 | 2403 | 107 | 0.27 | 28 | 41 |
| H2 | Substrate mix | 1.38 | 5.93±1.27 | 42.02±6.46 | 6.58 | 1560 | - | 7.18 | - | - |
| | Digestate | | 4.29±0.83 | 33.42±4.07 | 7.39 | 1998 | 69 | 0.85 | 18 | 39 |
| | Storage | | 2.86±0.17 | 23.25±2.00 | 7.32 | 1449 | - | 1.90 | na[h] | na[h] |
| L | Solid substrate | 1.82 | 77.04±0.55 | 85.69±2.09 | - | - | - | - | - | - |
| | Liquid substrate | | na[g] | na[g] | 7.72 | 1611 | - | 4.28 | - | - |
| | Digestate | | 9.23±0.17 | 51.46±1.89 | 7.46 | 1983 | 71 | 0.14 | 55 | 37 |
| | Storage | | 6.43±0.18 | 44.52±2.04 | 7.45 | 2565 | - | 0.11 | na[h] | na[h] |
| V | Substrate mix | 1.00 | 7.75±0.10 | 50.15±1.24 | 6.57 | 2061 | - | 11.49 | - | - |
| | Digestate 1 | | 5.15±1.65 | 30.70±3.60 | 7.69 | 3060 | 186 | 0.45 | 30 | 38 |
| | Digestate 2 | | 4.45±0.35 | 33.36±2.42 | 7.52 | 3090 | 130 | 0.67 | 30 | 38 |
| | Storage | | 1.92±0.10 | 16.17±2.37 | 7.74 | 2400 | - | 0.25 | na[h] | na[h] |
| K2 | liquid manure | 4.45 | 7.59±0.09 | 52.39±0.29 | 7.29 | 1704 | - | 5.50 | - | - |
| | Solid manure | | 35.81±0.30 | 74.78±1.12 | - | - | - | - | - | - |
| | Agnar | | 85.03±0.06 | 80.74±0.65 | - | - | - | - | - | - |
| | Digestate | | 5.55±0.56 | 38.03±3.30 | 8.01 | 2127 | 509 | 0.18 | 32 | 51 |
| | Storage 1 | | 5.04±0.49 | 37.16±2.63 | 7.66 | 2178 | - | 0.95 | 30 | na[h] |
| | Storage 2 | | 4.18±0.09 | 31.33±2.18 | 7.86 | 2307 | - | 1 | 30 | na[h] |
| B | Substrate mix | 3.11 | 9.16±0.10 | 58.18±1.57 | 6.55 | 2043 | - | 10.86 | - | - |
| | Digestate | | 7.33±0.07 | 49.87±0.83 | 6.87 | 2004 | 21 | 7.10 | 30 | 39 |
| | Storage | | 4.57±0.02 | 32.81±0.87 | 7.70 | 2187 | - | 0.49 | 21 | na[h] |
| J | Liquid substrate | 5.29 | 7.78±0.01 | 61.45±2.68 | 3.58 | 128 | - | 25.49 | - | - |
| | Solid substrate | | 38.79±0.89 | 78.28±1.32 | - | - | - | - | - | - |
| | Digestate 1 | | 32.69±0.46 | 68.83±2.52 | 7.30 | 1563 | 72 | 0.75 | 36 | 47 |
| | Digestate 2 | | 11.93±0.12 | 63.35±1.54 | 7.66 | 2412 | 240 | 0.32 | 36 | 47 |
| | Digestate 3 | | 13.06±0.07 | 62.09±1.26 | 7.66 | 2532 | 252 | 0.44 | 36 | 47 |
| | Liquid storage | | 11.91±0.11 | 61.12±1.91 | 7.67 | 2004 | - | 0.43 | 0 | na[h] |
| | Solid storage | | 6.22±0.02 | 44.53±3.10 | - | - | - | - | 10 | na[h] |
| K1 | Substrate mix | 6.68 | 9.40±0.16 | 52.27±3.04 | 7.14 | 1812 | - | 0.22 | - | - |
| | Digestate | | 7.06±0.15 | 46.37±1.07 | 7.82 | 2148 | 225 | 0.57 | 17 | 42 |
| | Storage | | 3.65±0.31 | 31.27±0.70 | 8.15 | 1473 | - | 0.33 | 70 | na[h] |

| | | | | | | | | | | |
|---|---|---|---|---|---|---|---|---|---|---|
| S | Substrate mix | 0.60 | 4.62±0.05 | 38.10±1.32 | 6.98 | 1458 | - | 5.35 | - | - |
| | Digestate | | 4.01±0.02 | 36.83±0.82 | 7.67 | 1776 | 89 | 0.08 | 29 | 35 |
| | Secondary digestate | | 3.94±0.01 | 31.59±0.67 | 7.48 | 1788 | 59 | 0.11 | 33 | 35 |
| N | Substrate mix | 0.78 | 5.43±0.17 | 40.15±1.93 | 6.99 | 1935 | - | 7.99 | - | - |
| | Digestate | | 3.14±0.02 | 27.79±1.24 | 7.80 | 1788 | 160 | 0.13 | 28 | 40 |
| | Storage | | 1.56±0.06 | 15.27±1.15 | 7.66 | 1863 | - | 0.16 | 21 | na[h] |

[a]Organic loading rate. [b]Volatile solids. [c]Total solids. [d]Free ammonia concentration calculated according to Calli et al. (2005). [e]Volatile fatty acids. [f]Hydraulic retention time. For digestate samples, this represents the hydraulic retention time of the biogas plant; for storage samples, it represents the storage duration prior to use as biofertiliser. [g]Not analysed due to insufficient sample quantity. [h]Information unavailable.

Table S3. Classification of antibiotic resistance gene (ARG) subtype and mobile genetic element (MGE) analyzed in this study.

| No. | ARG class | ARG subtype | ARG class | ARG subtype | MGE category | MGE element |
|---|---|---|---|---|---|---|
| 1 | MLSB | ermF_1 | Tetracycline | tetW | Integron | intI1_1 |
| 2 | MLSB | ermF | Tetracycline | tetM | Integron | intI3 |
| 3 | MLSB | lnuB | Tetracycline | tetQ | Integron | intI1_2 |
| 4 | MLSB | ermX_2 | Tetracycline | tet(44) | Integron | intI2 |
| 5 | MLSB | mel_1 | Tetracycline | tetD | Plasmid-associated | IncQ_oriT |
| 6 | MLSB | mphA | Tetracycline | tet(36)_1 | Plasmid-associated | IncI1_repI1 |
| 7 | MLSB | ermB_2 | Tetracycline | tetG | Plasmid-associated | IncP_oriT |
| 8 | MLSB | ermB_1 | Tetracycline | tetX | Plasmid-associated | IncW_trwAB |
| 9 | MLSB | ereA | Tetracycline | tetO_2 | Plasmid-associated | IncN_rep |
| 10 | MLSB | ermB_3 | Tetracycline | tetT | Plasmid-associated | trfA |
| 11 | MLSB | lnuC | Tetracycline | tetL_2 | Plasmid-associated | lncF_FIC |
| 12 | MLSB | ermE | Tetracycline | tetH | Plasmid-associated | IncHI2-smr0018 |
| 13 | MLSB | pncA | Tetracycline | tetR | Plasmid-associated | pAMBL |
| 14 | MLSB | mel | Tetracycline | tetB(P)_1 | Plasmid-associated | pAKD1 |
| 15 | MLSB | lnuF | Tetracycline | tet(32) | Plasmid-associated | IncN_korA |
| 16 | MLSB | ermA | Tetracycline | tetA_2 | Plasmid-associated | IncN_oriT |
| 17 | MLSB | ermC_2 | Tetracycline | tetS | Transposon-related | IS6/257 |
| 18 | MLSB | lnuA_1 | Tetracycline | tet(39) | Transposon-related | tnpA_6 |
| 19 | MLSB | erm(31) | Tetracycline | tetA/B_1 | Transposon-related | tnpA_7 |
| 20 | MLSB | vatB | Tetracycline | tetR_1 | Transposon-related | ISEcp1 |
| 21 | MLSB | ermO | Tetracycline | tetJ | Transposon-related | tnpA_5 |
| 22 | MLSB | ermY | Tetracycline | tetK | Transposon-related | IS1111 |
| 23 | MLSB | mefB | Tetracycline | tetE | Transposon-related | orf37-IS26 |
| 24 | MLSB | msrE | Tetracycline | tetA(P)_1 | Transposon-related | Tn5403 |
| 25 | MLSB | erm(35) | Tetracycline | tetC_2 | Transposon-related | IS256 |
| 26 | MLSB | erm(42) | Tetracycline | tet(38) | Transposon-related | IS1247_1 |
| 27 | MLSB | vatA | Vancomycin | vanA | Transposon-related | IS613 |
| 28 | MLSB | oleC | Vancomycin | vanTC_2 | Transposon-related | tnpA_1 |
| 29 | MLSB | ermX_1 | Vancomycin | vanYD_1 | Transposon-related | IS6100 |
| 30 | MLSB | msrA_1 | Vancomycin | vanHB | Transposon-related | IS630 |
| 31 | MLSB | lmrA_1 | Vancomycin | vanB_1 | Transposon-related | IS26_1 |
| 32 | MLSB | erm(36) | Vancomycin | vanC_5 | Transposon-related | tnpA_2 |
| 33 | MLSB | vatE_2 | Vancomycin | vanTE | Transposon-related | tnpA_3 |
| 34 | MLSB | ermD_2 | Vancomycin | vanYB | Transposon-related | IS1133 |
| 35 | MLSB | vgaB_1 | Vancomycin | vanRB | Transposon-related | IS1247_2 |
| 36 | MLSB | ermH | Vancomycin | vanSC_2 | Transposon-related | ISCR1 |
| 37 | MLSB | ermT_1 | Vancomycin | vanHD | Transposon-related | IS21-ISAs29 |
| 38 | MLSB | mphB | Vancomycin | vanRA_1 | Transposon-related | Tp614 |
| 39 | MLSB | erm(34) | Vancomycin | vanTG | Transposon-related | ISEfm1 |
| 40 | MLSB | ermD | Vancomycin | vanC_2 | Transposon-related | Tn3 |

| | | | | | | |
|---|---|---|---|---|---|---|
| 41 | MLSB | ermA_2 | Vancomycin | vanG | Transposon-related | IS3 |
| 42 | MLSB | vgaA_1 | Vancomycin | vanRC | Transposon-related | ISAba3 |
| 43 | MLSB | msrC_1 | Vancomycin | vanRC_2 | Transposon-related | Tn5 |
| 44 | MLSB | lsaC | Vancomycin | vanSB | Transposon-related | IS5/IS1182 |
| 45 | MLSB | ermD_1 | Vancomycin | vanXB | Transposon-related | IS200_2 |
| 46 | MLSB | vga(A)LC_1 | Vancomycin | vanWB | Transposon-related | tnpA_4 |
| 47 | MLSB | mel_3 | Vancomycin | vanRD | Transposon-related | ISPps |
| 48 | Quinolone | qepA | Vancomycin | vanD | Transposon-related | IS200_1 |
| 49 | Quinolone | qnrB_3 | Vancomycin | vanSA | Other-MGE | trbC |
| 50 | Quinolone | norA | Vancomycin | vanXA | Other-MGE | EAE_05855 |
| 51 | Quinolone | qnrB_2 | Trimethoprim | dfrA17_1 | Other-MGE | traN |
| 52 | Quinolone | qnrS2 | Trimethoprim | dfra17 | Other-MGE | cro |
| 53 | Quinolone | qnrD | Trimethoprim | dfrA1_1 | | |
| 54 | Quinolone | qnrB | Trimethoprim | dfrA27 | | |
| 55 | Quinolone | qnrVC1_VC3_VC6 | Trimethoprim | dfrA25 | | |
| 56 | Quinolone | qnrVC_2 | Trimethoprim | dfrB | | |
| 57 | Quinolone | qnrA | Trimethoprim | dfrA15 | | |
| 58 | Quinolone | qnrS_1 | Trimethoprim | dfrA22 | | |
| 59 | Beta Lactam | penA | Trimethoprim | dfrA12_1 | | |
| 60 | Beta Lactam | blaOXY | Trimethoprim | dfrG | | |
| 61 | Beta Lactam | blaSFO | Trimethoprim | dfrK | | |
| 62 | Beta Lactam | blaGOB | Trimethoprim | dfrA8 | | |
| 63 | Beta Lactam | cfxA | Trimethoprim | dfrA12 | | |
| 64 | Beta Lactam | blaOXY-1 | Trimethoprim | dfrA10 | | |
| 65 | Beta Lactam | blaMIR | Trimethoprim | dfrC | | |
| 66 | Beta Lactam | blaACT | Trimethoprim | dfrA19 | | |
| 67 | Beta Lactam | blaFOX | Trimethoprim | dfrAB4 | | |
| 68 | Beta Lactam | blaCTX-M | Aminoglycoside | aadA7 | | |
| 69 | Beta Lactam | blaSHV_4 | Aminoglycoside | aadA_4 | | |
| 70 | Beta Lactam | blaOXA-48 | Aminoglycoside | aadA_1 | | |
| 71 | Beta Lactam | blaNDM | Aminoglycoside | ANT(6)-Ia | | |
| 72 | Beta Lactam | cphA_1 | Aminoglycoside | aadA2_3 | | |
| 73 | Beta Lactam | blaDHA | Aminoglycoside | APH(6)-Ic | | |
| 74 | Beta Lactam | blaOCH | Aminoglycoside | APH(7")-Ib | | |
| 75 | Beta Lactam | blaCMY_2 | Aminoglycoside | AAC(3)-IIa_IId | | |
| 76 | Beta Lactam | blaCARB | Aminoglycoside | APH(3')-Ib | | |
| 77 | Beta Lactam | blaACC | Aminoglycoside | APH(6)-Id | | |
| 78 | Beta Lactam | blaROB | Aminoglycoside | ANT(6)-Ib | | |
| 79 | Beta Lactam | ccrA | Aminoglycoside | AAC(3)-III | | |
| 80 | Beta Lactam | cfiA | Aminoglycoside | APH(9)-Ib | | |
| 81 | Beta Lactam | pbp5 | Aminoglycoside | APH(6)-Ia | | |
| 82 | Beta Lactam | blaOXA-51 | Aminoglycoside | aadA16 | | |
| 83 | Beta Lactam | blaCTX-M-5 | Aminoglycoside | ANT(6)-Ia_1 | | |

| | | | | |
|---|---|---|---|---|
| 84 | Beta Lactam | blaPER | Aminoglycoside | AAC(3)-Iva |
| 85 | Beta Lactam | blaMOX/blaCMY | Aminoglycoside | ANT(2")-Ia |
| 86 | Beta Lactam | pbp | Aminoglycoside | aadA6 |
| 87 | Beta Lactam | blaL1 | Aminoglycoside | rmtB |
| 88 | Beta Lactam | imiR | Aminoglycoside | APH(3')-Ia |
| 89 | Beta Lactam | blaPDC | Aminoglycoside | AAC(3)-Ib |
| 90 | Beta Lactam | blaVIM | Aminoglycoside | AAC(6')-Ir |
| 91 | Beta Lactam | blaSME | Aminoglycoside | aadA2_1 |
| 92 | Beta Lactam | blaTEM | Aminoglycoside | APH(3')-IIIa_1 |
| 93 | Beta Lactam | blaADC | Aminoglycoside | APH(3')-Ia |
| 94 | Beta Lactam | blaBEL | Aminoglycoside | AAC(3)-Xa |
| 95 | Beta Lactam | blaKPC | Aminoglycoside | AAC(3)-IVa_1 |
| 96 | Beta Lactam | blaHERA | Aminoglycoside | AAC(6')-IIc |
| 97 | Beta Lactam | blaACC-1 | Aminoglycoside | ANT(4')-Ib |
| 98 | Beta Lactam | blaSST | Aminoglycoside | apmA |
| 99 | Beta Lactam | blaIMI | Aminoglycoside | aadA10 |
| 100 | Beta Lactam | blaCARB_1 | Aminoglycoside | aadA9_1 |
| 101 | Beta Lactam | cepA | Aminoglycoside | AAC(6')-Iz |
| 102 | Beta Lactam | blaLEN | Aminoglycoside | APH(7")-Ia |
| 103 | Beta Lactam | mecA | Aminoglycoside | AAC(3)-Id |
| 104 | Beta Lactam | cphA_3 | Aminoglycoside | AAC(6')-Ib_1 |
| 105 | Beta Lactam | blaZ | Aminoglycoside | ANT(4') |
| 106 | Beta Lactam | blaVEB | Aminoglycoside | APH(3")-Ib |
| 107 | Beta Lactam | blaCTX-M-8 | Aminoglycoside | AAC(6')-Iv_Ih |
| 108 | Beta Lactam | blaB | Aminoglycoside | APH(3')-VIIIa |
| 109 | Beta Lactam | blaIND | Aminoglycoside | AAC(6')-Ig |
| 110 | Beta Lactam | blaGES | Aminoglycoside | APH(3')-VIa |
| 111 | Beta Lactam | bla1 | Aminoglycoside | aadA5_2 |
| 112 | Beta Lactam | blaTLA | Aminoglycoside | AAC(6')-Ie-APH(2")-Ia |
| 113 | MDR | oprD | Aminoglycoside | AAC(6')-Is_Iu_Ix |
| 114 | MDR | mdtA | Aminoglycoside | AAC(6')-Iy |
| 115 | MDR | mdtH | Aminoglycoside | AAC(6')-Ie-APH(2")-Ia_1 |
| 116 | MDR | tcrB | Aminoglycoside | AAC(6')-II |
| 117 | MDR | emrD_1 | Aminoglycoside | armA_1 |
| 118 | MDR | qacEdelta1_4 | Aminoglycoside | AAC(3)-IIc |
| 119 | MDR | mepA | Aminoglycoside | APH(9)-Ia_1 |
| 120 | MDR | arsA | Aminoglycoside | AAC(6')-I |
| 121 | MDR | pbrT | Aminoglycoside | APH(3')-VIIa |
| 122 | MDR | cadC | Aminoglycoside | AAC(6')-Im |
| 123 | MDR | sugE | Aminoglycoside | AAC(6')-Ij |
| 124 | MDR | czcA | Aminoglycoside | APH(2")-Iia |
| 125 | MDR | mexB | Aminoglycoside | AAC(6')-I-43 |
| 126 | MDR | ttgA | Aminoglycoside | AAC(6')-Iw |

| | | | | |
|---|---|---|---|---|
| 127 | MDR | mdtE | Aminoglycoside | armA_2 |
| 128 | MDR | tolC_2 | Sulfonamide | sul2_2 |
| 129 | MDR | acrF | Sulfonamide | sul1_2 |
| 130 | MDR | qacA/B | Sulfonamide | sul4 |
| 131 | MDR | pcoA | Sulfonamide | sul3_1 |
| 132 | MDR | acrR_1 | Sulfonamide | folA_1 |
| 133 | MDR | copA | Sulfonamide | folP_2 |
| 134 | MDR | acrA_1 | Phenicol | cmlV |
| 135 | MDR | acrB_1 | Phenicol | catQ |
| 136 | MDR | qacH_1 | Phenicol | cmlA_2 |
| 137 | MDR | oqxA | Phenicol | cmlA_4 |
| 138 | MDR | mdtG_1 | Phenicol | cmxA |
| 139 | MDR | terW | Phenicol | floR_1 |
| 140 | MDR | adeA | Phenicol | ceoA |
| 141 | MDR | cmr | Phenicol | mdtL_1 |
| 142 | MDR | mexE | Phenicol | floR |
| 143 | MDR | mexA | Phenicol | mdtL |
| 144 | MDR | adeI | Phenicol | catIII |
| 145 | MDR | mdsA | Phenicol | catB9 |
| 146 | MDR | marR_3 | Phenicol | catB3 |
| 147 | MDR | bexA_1 | Phenicol | cat |
| 148 | MDR | pmrA | Phenicol | optrA |
| 149 | MDR | mtrE | Phenicol | catB8 |
| 150 | MDR | emrB/qacA_1 | Phenicol | catP |
| 151 | MDR | cfr | Phenicol | catA1 |
| 152 | Other | bacA | Phenicol | cat(pC221) |
| 153 | Other | qacEdelta1_3 | Phenicol | catB2 |
| 154 | Other | qacEdelta1_1 | Phenicol | catA2 |
| 155 | Other | merA | Phenicol | fexA |
| 156 | Other | SAT-4 | Other | nimE |
| 157 | Other | MCR-1 | Other | fosX |
| 158 | Other | fabK | Other | fosB |
| 159 | Other | arr-3 | Other | MCR-2 |
| 160 | Other | nisB_1 | Other | crAss56 |
| 161 | Other | arr-2 | Other | crAss64 |
| 162 | Other | ttgB | | |

Table S4. Baseline relative abundance and directional shifts of mobile genetic elements (MGEs).

| MGE category | No. of subtypes | Baseline relative abundance (median) | Baseline IQR | Median Δ (digester − substrate) | Δ / baseline (%) | p_ adj_ fdr | Directional support |
|---|---|---|---|---|---|---|---|
| Integrons | 4 | $1.56 \times 10^{-3}$ | $1.38 \times 10^{-2}$ | $-3.76 \times 10^{-4}$ | -24.1 | 0.490 | No |
| Plasmid-associated | 12 | $2.92 \times 10^{-5}$ | $7.19 \times 10^{-5}$ | $8.76 \times 10^{-7}$ | 3.0 | 0.930 | No |
| Transposon-related | 32 | $6.75 \times 10^{-4}$ | $3.63 \times 10^{-3}$ | $-2.19 \times 10^{-4}$ | -32.5 | 0.008 | Yes |
| Other-MGE | 4 | $1.01 \times 10^{-4}$ | $3.87 \times 10^{-4}$ | $-1.06 \times 10^{-5}$ | -10.6 | 0.685 | No |
| Overall MGEs (pooled) | 52 | $4.68 \times 10^{-4}$ | $2.81 \times 10^{-3}$ | $-1.19 \times 10^{-4}$ | -25.4 | 0.006 | Yes |

Baseline relative abundance was calculated as the median and interquartile range (IQR) across substrate mix samples. Median Δ values represent digester-level directional shifts in relative abundance (digester − substrate). Relative changes (Δ/baseline, %) are provided for contextual interpretation only and were not subjected to additional statistical testing. Directional support indicates whether the digester-level median Δ deviated from zero based on Wilcoxon signed-rank tests with FDR correction.

Table S5. Top hub nodes identified across different co-occurrence networks

| Network type | Node name | Node type | Degree |
|---|---|---|---|
| Genus-ARG | ermF_1 | ARG | 12 |
| Genus-ARG | AAC(3)-III | ARG | 8 |
| Genus-ARG | ANT(6)-Ia_1 | ARG | 8 |
| Genus-ARG | tcrB | ARG | 6 |
| Genus-ARG | lnuB | ARG | 5 |
| Genus-ARG | sugE | ARG | 5 |
| Genus-ARG | ANT(6)-Ia | ARG | 4 |
| Genus-ARG | tet(36)_1 | ARG | 3 |
| Genus-ARG | tet(44) | ARG | 3 |
| Genus-ARG | ermF | ARG | 3 |
| Genus-ARG | Acholeplasma | Genus | 13 |
| Genus-ARG | Syntrophomonas | Genus | 8 |
| Genus-ARG | HN-HF0106 | Genus | 6 |
| Genus-ARG | Ruminiclostridium | Genus | 6 |
| Genus-ARG | Sedimentibacter | Genus | 5 |
| Genus-ARG | Dethiobacter | Genus | 5 |
| Genus-ARG | Caldicoprobacter | Genus | 4 |
| Genus-ARG | UCG-004 | Genus | 4 |
| Genus-ARG | Christensenellaceae R-7 group | Genus | 3 |
| Genus-ARG | HT002 | Genus | 3 |
| Genus-MGE | Bacteroides | Genus | 4 |
| Genus-MGE | Bifidobacterium | Genus | 3 |
| Genus-MGE | Peptostreptococcus | Genus | 2 |
| Genus-MGE | Caldicoprobacter | Genus | 1 |
| Genus-MGE | Lentimicrobium | Genus | 1 |
| Genus-MGE | Corynebacterium | Genus | 1 |
| Genus-MGE | Fastidiosipila | Genus | 1 |
| Genus-MGE | Christensenellaceae R-7 group | Genus | 1 |
| Genus-MGE | Atopostipes | Genus | 1 |
| Genus-MGE | UCG-005 | Genus | 1 |
| Genus-MGE | tnpA_7 | MGE | 6 |
| Genus-MGE | trbC | MGE | 3 |
| Genus-MGE | IS6/257 | MGE | 2 |
| Genus-MGE | IncQ_oriT | MGE | 2 |
| Genus-MGE | tnpA_5 | MGE | 1 |
| Genus-MGE | IS256 | MGE | 1 |
| Genus-MGE | IS1111 | MGE | 1 |
| Genus-MGE | IS630 | MGE | 1 |
| Genus-MGE | Tp614 | MGE | 1 |
| Genus-MGE | Tn5 | MGE | 1 |

| | | | |
|---|---|---|---|
| ARG-MGE | tetO_2 | ARG | 15 |
| ARG-MGE | blaOXY | ARG | 13 |
| ARG-MGE | APH(3')-Ib | ARG | 11 |
| ARG-MGE | oprD | ARG | 8 |
| ARG-MGE | rmtB | ARG | 8 |
| ARG-MGE | penA | ARG | 7 |
| ARG-MGE | tetD | ARG | 7 |
| ARG-MGE | ereA | ARG | 7 |
| ARG-MGE | cfxA | ARG | 7 |
| ARG-MGE | blaMIR | ARG | 7 |
| ARG-MGE | orf37-IS26 | MGE | 29 |
| ARG-MGE | ISEcp1 | MGE | 21 |
| ARG-MGE | IS21-ISAs29 | MGE | 20 |
| ARG-MGE | IS630 | MGE | 19 |
| ARG-MGE | tnpA_5 | MGE | 18 |
| ARG-MGE | ISCR1 | MGE | 16 |
| ARG-MGE | Tn5403 | MGE | 16 |
| ARG-MGE | IS1111 | MGE | 15 |
| ARG-MGE | EAE_05855 | MGE | 13 |
| ARG-MGE | intI3 | MGE | 11 |

Top hub nodes identified in the co-occurrence networks among bacterial genera, antibiotic resistance genes (ARGs), and mobile genetic elements (MGEs) across 42 samples. Co-occurrence networks were constructed based on Spearman's rank correlation analysis, and only significant correlations ($|\rho| > 0.7$ and false discovery rate (FDR)–adjusted $p < 0.05$) were retained. Hub nodes were defined based on degree values, representing the number of significant correlations with other nodes within each network.

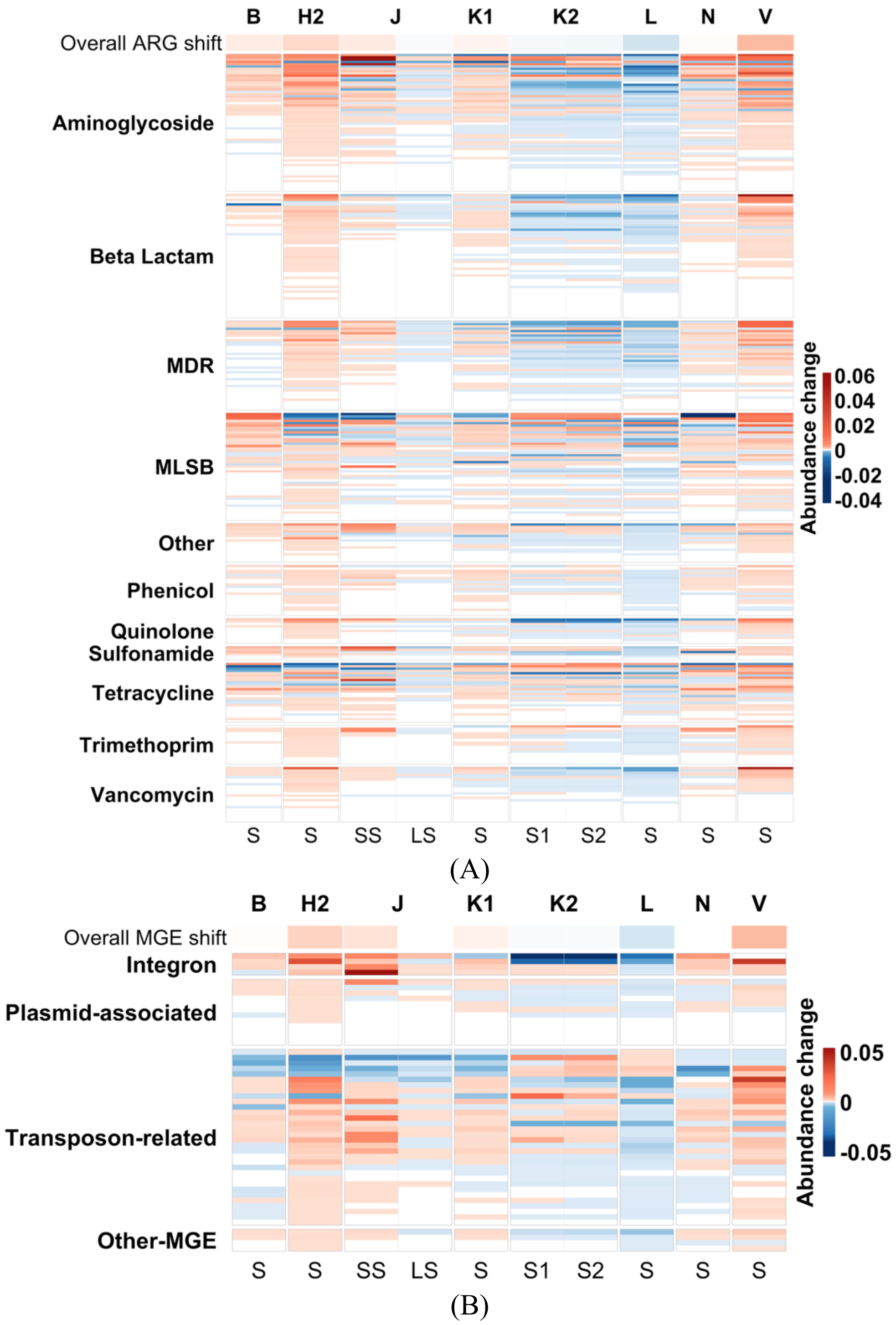


Fig. S1. Fate of antibiotic resistance genes (ARGs) and mobile genetic elements (MGEs) during storage in full-scale biogas plants. Changes in relative abundance are expressed as the difference between storage digestate and the corresponding digester samples (storage − digester). Red and blue indicate increases and decreases in relative abundance, respectively. (A) ARG subtypes grouped by antibiotic classes. (B) MGEs grouped by functional categories. Columns represent individual storage systems; S, SS, and LS denote storage, solid storage, and liquid storage, respectively. For both panels, the top bar summarizes the overall median change (Δ) in relative abundance for each storage system, with blue and red indicating negative and positive deviations, respectively. The colour scale of the top bar is independent of that of the main heatmap and is used solely to indicate the direction of overall change.

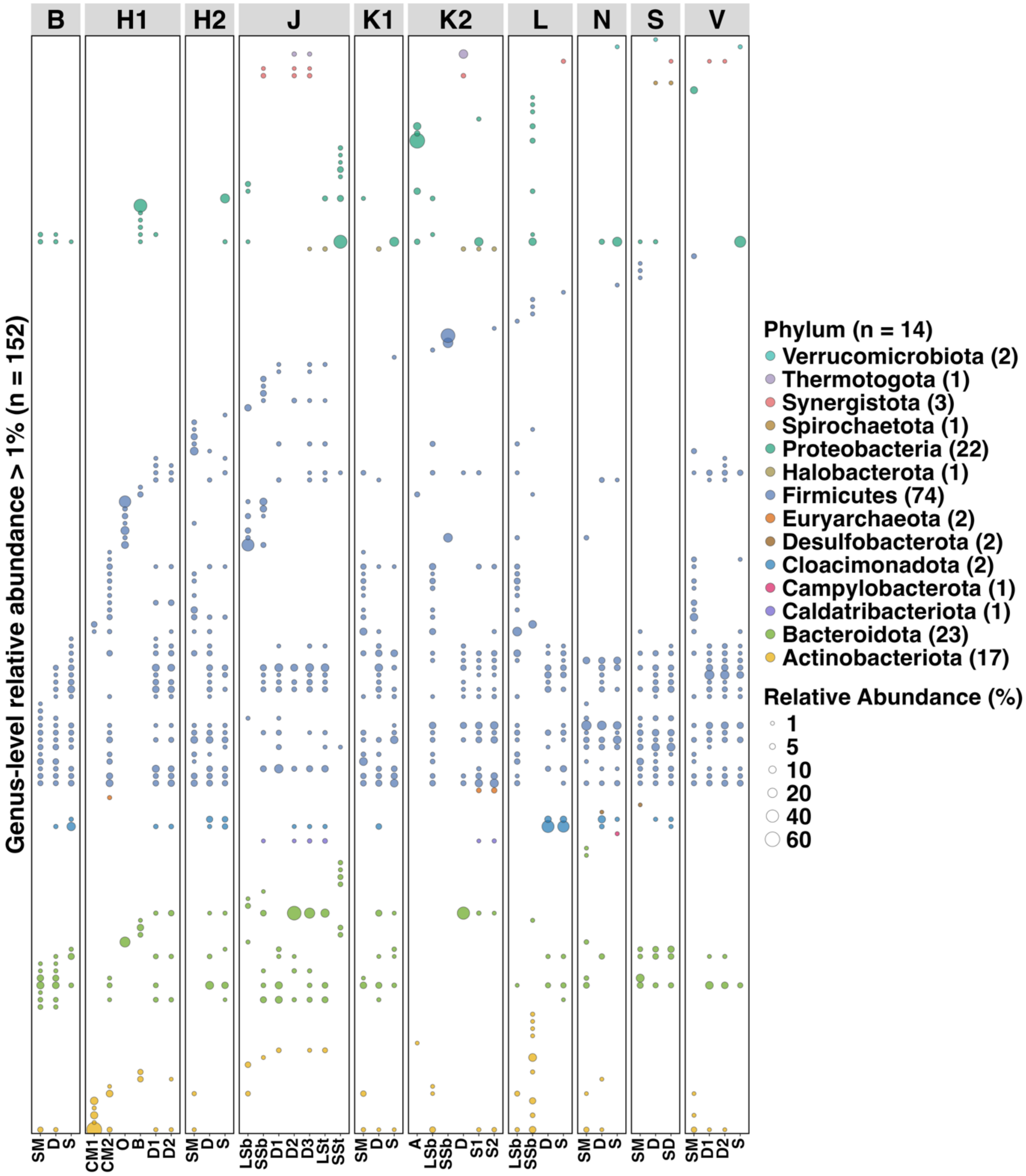


Fig. S2. Genus-level bacterial community composition across substrates, digestates, and storage samples from the ten investigate biogas plants. Bubble plot showing bacterial genera with relative abundance > 1% across all samples, comprising a total of 152 genera. Bubble size corresponds to genus-level relative abundance (%), while colours indicate bacterial phyla. Numbers in parentheses denote the total number of genera detected within each phylum across all samples. Abbreviations: SM, substrate mix; D, D1, D2, digestate samples; SD, secondary digestate; S, storage sample; CM1, chicken manure; CM2, cow manure; O, oat water; B, bedding material; LSb, liquid substrate; SSb, solid substrate; A, chaff; LSt, liquid storage; SSt, solid storage.

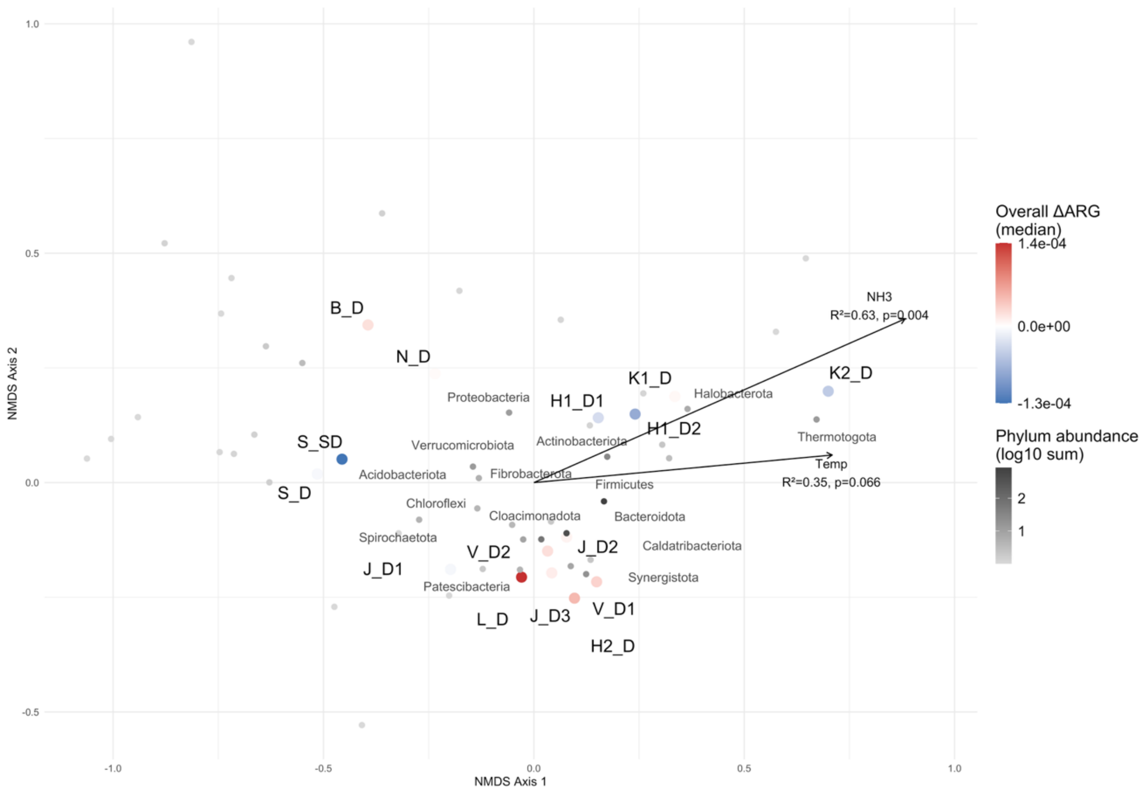


Fig. S3. Non-metric multidimensional scaling (NMDS) ordination of full-scale biogas digesters based on microbial community composition at the phylum level. Digester samples are coloured according to the overall median shift in ARG relative abundance (ΔARG), calculated as the difference between digesters and their corresponding substrates (Digester − Substrate). Colour gradients range from blue (negative change) to red (positive change), with white indicating no net change. Arrows represent fitted physicochemical parameters showing significant correlations with the ordination ($p < 0.1$). Grey-scale shading of phylum labels indicates summed phylum-level abundance ($log_{10}$-transformed) across all digesters; only the 15 most abundant phyla are labelled to improve visual clarity.
